# A multi-task deep-learning system for predicting membrane associations and secondary structures of proteins

**DOI:** 10.1101/2020.12.02.409045

**Authors:** Bian Li, Jeffrey Mendenhall, John A. Capra, Jens Meiler

**Affiliations:** Department of Chemistry and Center for Structural Biology, Vanderbilt University, Nashville, TN 37203, USA; Department of Biological Sciences, Vanderbilt University, Nashville, TN 37203, USA; Bakar Computational Health Sciences Institute and Department of Epidemiology and Biostatistics, University of California, San Francisco, CA 94143, USA; Institute for Drug discovery, University Leipzig Medical School, Leipzig, Germany

**Keywords:** secondary structure prediction, membrane topology prediction, multi-task deep learning

## Abstract

Accurate prediction of secondary structures and transmembrane segments is often the first step towards modeling the tertiary structure of a protein. Existing methods are either specialized in one class of proteins or developed to predict one type of 1D structural attributes (secondary structure, topology, or transmembrane segment). In this work, we develop a new method for simultaneous prediction of secondary structure, transmembrane segment, and transmembrane topology with no *a priori* assumption on the class of the input protein sequence. The new method, Membrane Association and Secondary Structures of Proteins (MASSP) predictor, uses multi-tiered neural networks that incorporate recent innovations in machine learning. The first tier is a multi-task multi-layer convolutional neural network (CNN) that learns patterns in image-like input position-specific-scoring matrices (PSSMs) and predicts residue-level 1D structural attributes. The second tier is a long short-term memory (LSTM) neural network that treats the predictions of the first tier from the perspective of natural language processing and predicts the class of the input protein sequence. We curated a non-redundant data set consisting of 54 bitopic, 241 multi-spanning TM-alpha, 77 TM-beta, and 372 soluble proteins, respectively for training and testing MASSP. For secondary structure prediction, the mean three-state accuracy (Q3) of MASSP is 0.830, better than the Q3 of PSIPRED (0.829) and that of SPINE-X (0.813) and substantially better than that of Jufo9D (0.762) and RaptorX-Property (0.741). The mean segment overlap score (SOV) of MASSP is 0.752, gaining at least 7.7% improvement over all the other four methods. For transmembrane topology prediction, MASSP has a performance comparable to OCTOPUS and substantially better than MEMSAT3 and TMHMM2 on TM-alpha proteins, and on TM-beta proteins, MASSP is significantly better than both BOCTOPUS2 and PRED-TMBB2. By integrating prediction of secondary structure and transmembrane segments in a deep-learning framework, MASSP improves performance over previous methods, has broader applicability, and enables proteome scale predictions.

## Introduction

Proteins are molecular machines that drive most biochemical processes in the cell. As the tertiary structure of a protein largely determines its molecular function, the elucidation of a protein’s structure is essential to our understanding of the function of the protein, its dysfunction in diseases, and the design of therapeutic molecules to target the protein. Although several experimental techniques exist for protein tertiary structure determination, determining the structures of all proteins and disease-associated variants by experimental techniques remains impractical. Thus, computational prediction of protein tertiary structures from amino acid sequences is an area of active research (Li et al., 2018). While exciting progress was made in the past few years, protein tertiary structure prediction is still an unsolved problem (Senior et al., 2020; Shrestha et al., 2019).

Accurate prediction of secondary structure is often the first step towards predicting tertiary structure. Over the past decades, a plethora of computational methods were developed for secondary structure prediction. Some of the first methods relied on statistical propensities of amino acids to form particular structures (Chou and Fasman, 1974), simple nearest-neighbor algorithms that involve in finding short sequences of known structure from databases that closely match stretches of the query sequence (Frishman and Argos, 1997), or explicit modeling of the contribution of neighboring residues make to the probability of a given structure state (Garnier et al., 1996). The accuracy of these methods hovered under 70% for the so-called Q3, a three-state prediction that classifies amino acids in likely helix (H), strand (extended, E), or coil (C) states. More accurate methods, better than 70% in Q3, were developed that use machine-learning-based models (Rost and Sander, 1993b). The machine-learning models powering these methods are often artificial neural networks (Jones, 1999; Meiler and Baker, 2003; Meiler et al., 2001; Qian and Sejnowski, 1988; Rost, 1996; Rost and Sander, 1993a, b) that take sequence profiles generated from multiple sequence alignment as input (Jones, 1999; Rost, 1996) or couple sequence profiles with nonlocal interactions from predicted tertiary structures to boost the accuracy of secondary structure prediction (Meiler and Baker, 2003). Recently, motivated by unprecedented performance of deep neural networks in computer vision problems (LeCun et al., 2015), deeper neural network architectures, such as convolutional neural networks (CNNs) and recurrent neural networks were applied to protein secondary structure prediction with accuracy improving to above 80% in Q3 (Guo et al., 2019; Torrisi et al., 2020; Wang et al., 2016b; Zhang et al., 2018). The reader is referred to (Torrisi et al., 2020; Yang et al., 2018) for a broad review of the history and state of the art of secondary structure prediction.

For integral membrane proteins (IMPs), an additional essential component of predicting the tertiary structure is to first locate all the transmembrane segments and the orientation of the protein with respect to the membrane. These tasks are collectively known as “topology prediction”. Several algorithms have been developed over the past decades for predicting the topology of either α-helical or β-barrel IMPs. The earliest method of Kyte and Doolittle (Kyte and Doolittle, 1982) for transmembrane helix (TMH) prediction computes simple “hydropathy plots” to identify probable transmembrane segments. While this method is conceptually simple and easy to implement, it can neither predict the “inside-outside phasing” of the helices relative to the cytoplasm, i.e. topology, nor the location of transmembrane strands for β-barrels. By combining hydropathy analysis and the “positive-inside rule”, the observation that positively charged residues are more abundant in cytoplasmic as compared to periplasmic regions of bacterial inner membrane proteins (von Heijne, 1986), a method dubbed TOP-PRED was developed for predicting TMHs and their topologies (von Heijne, 1992). While TOP-PRED identified all 135 TMHs in the 24 proteins tested with only one overprediction and correctly predicted the topology of 22 proteins, no results were reported as to how accurate TOP-PRED can locate the boundaries (N- and C-terminal ends) of TMHs. Aimed at predicting both the boundaries and topology of TMHs, the PHDhtm method, whose driving predictor is a two-level neural network system trained on 69 transmembrane proteins, was developed (Rost et al., 1995; Rost et al., 1996). TMHMM and HMMTOP (Tusnady and Simon, 1998) are the first methods to model membrane topology of IMPs with hidden Markov models. The state-of-the-art methods for membrane region detection are MEMSAT3 (Jones, 2007) and OCTOPUS (Viklund and Elofsson, 2008) (for TM-alpha proteins) and BOCTOPUS2 (Hayat et al., 2016) and PRED-TMBB2 (Tsirigos et al., 2016) (for TM-beta proteins). However, these methods identify transmembrane regions in 4-10% of globular proteins (depending on the data set), a rate which is unacceptable for genomic annotation and sub-optimal for use in tertiary structure prediction. The interested reader is referred to (Fleishman and BenTal, 2006; Koehler Leman et al., 2015) for a more extensive review on the topic.

While these existing methods have been widely used, they are either specialized in one class of proteins or developed to predict one type of structural features (secondary structure, topology, or transmembrane segment). Previously, our group developed a method called Jufo9D that simultaneously predicts secondary structures and transmembrane spans (Leman et al., 2013). However, Jufo9D lacked the essential capability of predicting transmembrane topology. Further, Jufo9D displayed a somewhat reduced accuracy when tested in the 11^th^ community-wide Critical Assessment of protein Structure Prediction experiment (Fischer et al., 2016). In the current work, we leverage recent advances in deep neural network (DNN) architectures and develop a novel method that simultaneous predicts secondary structures, and for membrane proteins, the location and topology of transmembrane segments. The method is termed Membrane Association and Secondary Structure of Proteins (MASSP) predictor. MASSP uses multi-tiered DNNs that incorporate recent innovations in machine learning. The first tier is a multi-task multi-layer CNN that predicts residue-level structural features. The second tier is a long short-term memory (LSTM) neural network that treats the predictions of the first tier from the perspective of natural language processing and predicts the protein class of the input amino acid sequence. MASSP has a broader application domain than previous methods, because it simultaneously predicts secondary structure, membrane association, membrane topology for bitopic, multi-spanning TM-alpha, and TM-beta integral membrane proteins. The preliminary version of this method was successfully adopted in two previous studies and was favored over Jufo9D for predicting TMHs for *de novo* tertiary structure prediction of alpha-helical transmembrane (TM-alpha) proteins (Fischer et al., 2016; Li et al., 2017). In this manuscript, we describe the method in detail, demonstrate its broader applicability, and discuss aspects of its improved performance compared to some of the most popular methods in the field.

## Results

### Overview of MASSP

MASSP starts without an *a priori* assumption about the class of the input protein. It was designed as a two-tier prediction system to work with both soluble and membrane proteins (Fig. 1). The first tier is a multi-task multi-layer CNN that predicts residue-level structural features (e.g. secondary structure types, membrane association and topology). The second tier is an LSTM neural network that treats the predictions of the first tier as natural language input and predicts the protein class of the input amino acid sequence, i.e. bitopic, TM-alpha, beta barrel (TM-beta), or soluble.

**Figure 1.**
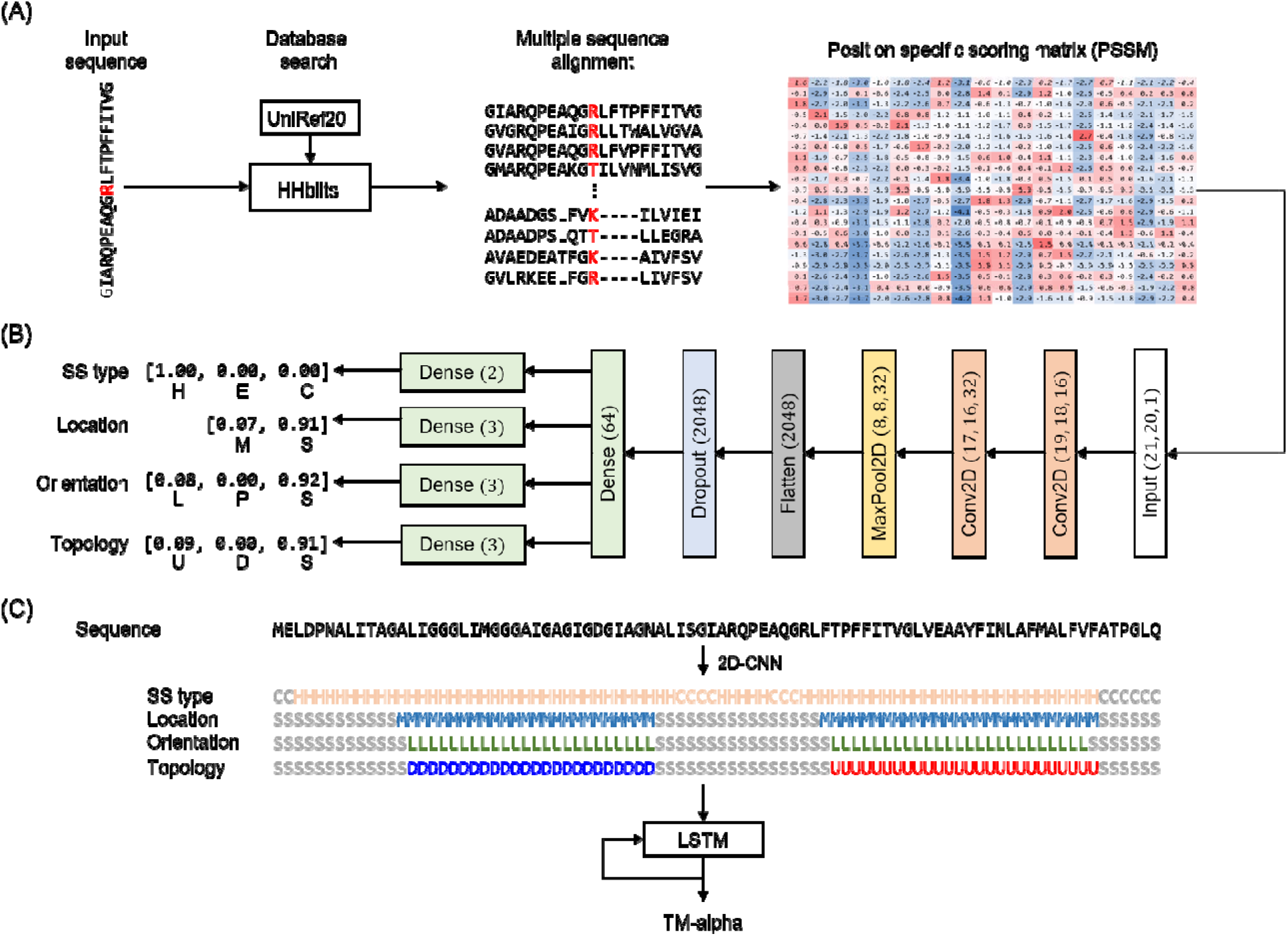
Design of the MASSP framework. MASSP is a hierarchical prediction system with two predictors. The first predictor is a multi-layer 2D convolutional neural network-based (2D-CNN) residue-level classifier trained to predict structural attributes of each residue in the input protein sequence (A and B). MASSP predicts four categories of residue-level attributes, namely secondary structure types (helix, strand, or coil, indicated by H, S, or C), location (membrane or solution, indicated by M or S), orientation (lipid-, pore-facing, or soluble indicated by L, P, or S), and transmembrane topology (up, down, or soluble indicated by U, D, or S). The second predictor is a long short-term memory (LSTM) recurrent neural network-based sequence-level classifier trained to predict the protein class (bitopic, TM-alpha, TM-beta, soluble) of the input amino acid sequence (C).

Given an amino acid sequence, MASSP calls HHblits (version 3.0) (Remmert et al., 2012) to search against the UniRef20 sequence database for homologous proteins and to create a multiple sequence alignment (MSA) of the hits to the input sequence. The MSA contains essential evolutionary information and is then used as input to an in-house C++ utility to compute the corresponding position-specific scoring matrix (PSSM, see Methods). To predict the target attributes of a given residue, MASSP takes as input a 21 × 20 matrix consisting of a sliding window of 21 positions around the residue of interest (Fig. 1A). The core algorithm of MASSP is a multi-output deep 2D CNN that simultaneously predicts the secondary structure type (helix, strand, or coil, indicated by H, E, or C in Fig. 1B), location (membrane or solution, indicated by M or S in Fig. 1B), orientation (lipid-, pore-facing, or soluble indicated by L, P, or S in Fig. 1B), and topology of the residue of interest (up, down, or soluble indicated by U, D, or S in Fig. 1B). While a separate model could be trained for each of the target attributes, we chose to learn to jointly predict all target attributes simultaneously, because these attributes aren’t necessarily independent. Once the 1D structural attributes of the input protein sequence are predicted by the first-tier CNN, the predicted symbols for each residue are combined to form “words”, which are then concatenated to form a “sentence”. The “semantic” meaning of this sentence (i.e. the class of the input protein sequence) is classified using an LSTM network (Fig. 1C).

We tuned the architecture of MASSP by following a workflow of developing deep-learning models recommended in (Chollet, 2018) (see Methods). For the CNN, we started out with a model that has a total of 1339 parameters. This “small” model confirmed that patterns in the input matrix are learnable (Fig. S1A). We then scaled up the model to 198187 parameters, which is sufficiently powerful to the point that overfitting is obvious (Fig. S1B). We continued several rounds of hyperparameter tuning by scaling down the size of the architecture and adding regularization (see Fig. S1C and S1D for two examples). The architecture of LSTM was similarly tuned. In the end, we selected a model with 136,651 parameters for training, shown in Fig. 1B, that has the optimal overall prediction accuracy on the independent validation set.

We developed MASSP using the experimental structures of 241 TM-alpha, 54 bitopic, and 77 TM-beta proteins. This data set was further augmented by adding 372 (accounting for 50% of the dataset) soluble protein subunits homology-reduced at 25% pairwise sequence identity level. The dataset was split according to the ratio 8:1:1 as training:validation:test (Tables S1, S2, S3). We used the validation set to tune hyperparameters of MASSP, such as number of layers and number of neurons in each layer, of the neural networks, and size of the input matrix (see Methods). The final selected architecture of MASSP, as described above, was tested on the test set that consists of five bitopic, 24 TM-alpha, eight TM-beta, and 37 soluble proteins, contributing a total of 18571 residues for which all four target attributes are available. The performance of MASSP was evaluated using the Q3 and the fractional segment overlap (SOV) measures for prediction accuracy. Q3 is the percentage of correctly predicted residues on all three types secondary structure, transmembrane topology, or in the case of TM-beta proteins, residue orientation. The SOV measure counts the fractional extent to which predicted and experimental segments of secondary structure or transmembrane region overlap, with some allowance for non-matching residues at the ends (see Methods) (Rost et al., 1994; Zemla et al., 1999).

### Dataset statistics

We split the data set according to the ratio 8:1:1 as training:validation:test while maintaining the constant fractions of each class of proteins in each subset. As shown in Fig. 2A, each of the three secondary structure types is evenly represented in the three subsets for TM-alpha, TM-beta, and soluble proteins. Some biases exist for bitopic proteins because a single bitopic protein with a large soluble domain could easily skew their representation among the three subsets. Likewise, membrane topologies are also similarly represented in the three subsets for all integral membrane protein classes (Fig. 2B). Soluble proteins are all considered to have a topology of outside the membrane. These three subsets each have 165080, 18293, 18571 training examples (residues) for which all four target attributes are available. For TM-alpha proteins, there are 1481, 150, and 158 transmembrane helices in the training, validation, and test sets respectively, and for TM-beta proteins, there are 878, 94, and 102 membrane-spanning beta strands, respectively. A breakdown of residue counts according to secondary structure types, location, transmembrane topology and be found in Table S4.

**Figure 2.**
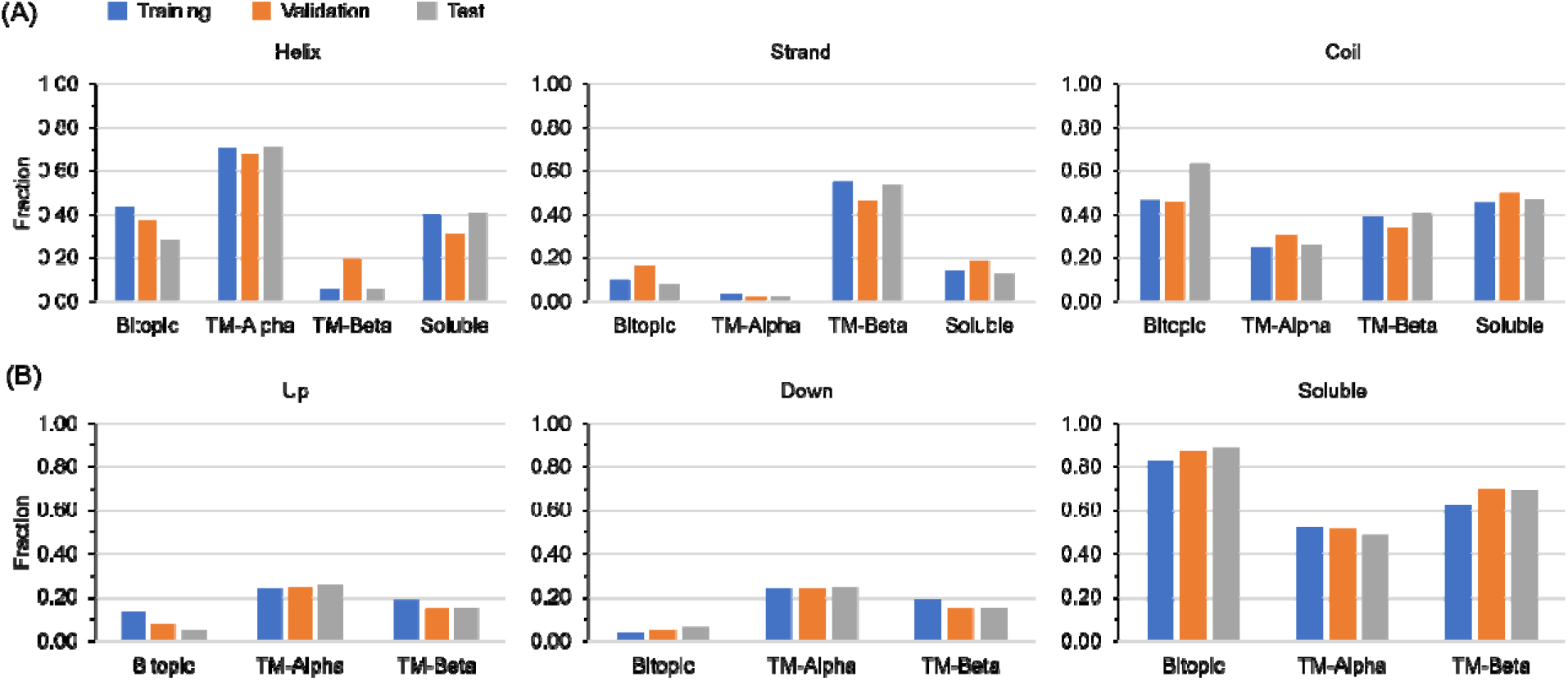
Residue secondary structures and topologies are evenly partitioned across protein types for training and evaluation. (A) The distribution of secondary structure types among training, validation, and test sets for all four protein classes. (B) The distribution of residue topology types among training, validation, and test sets for three protein classes. The distribution for soluble proteins is omitted because the topology types of all soluble protein residues are labeled as “Out”.

### MASSP accurately predicts structural attributes

Overall, MASSP achieves accurate simultaneous prediction of secondary structure, residue orientation, and transmembrane topology, with Q3s of 0.832, 0.942, and 0.922, respectively (Fig. 3A). When the overall accuracy was decomposed according to protein classes, MASSP achieved the highest Q3 for secondary structure prediction for TM-alpha proteins (0.874) and the lowest for soluble proteins (0.816) (Fig. 3B and Table S5). This is likely because the lipid bilayer imposes several constraints on structures of IMPs, simplifying their secondary structure prediction. Remarkably, for the three membrane related attributes, MASSP rarely predicts residues of soluble proteins to be membrane associated, as manifested by the near perfect accuracy in predicting location, orientation, and topology of residues of soluble proteins (Fig. 3B). Among the three classes of membrane associated proteins, the three membrane related attributes are much easier to predict for both bitopic and TM-beta proteins than for TM-alpha proteins. This is likely because the folds of TM-alpha proteins are generally much more complex than bitopic or TM-beta proteins. Contingency matrices for each of the structural attribute prediction tasks can be found in Table S6.

**Figure 3.**
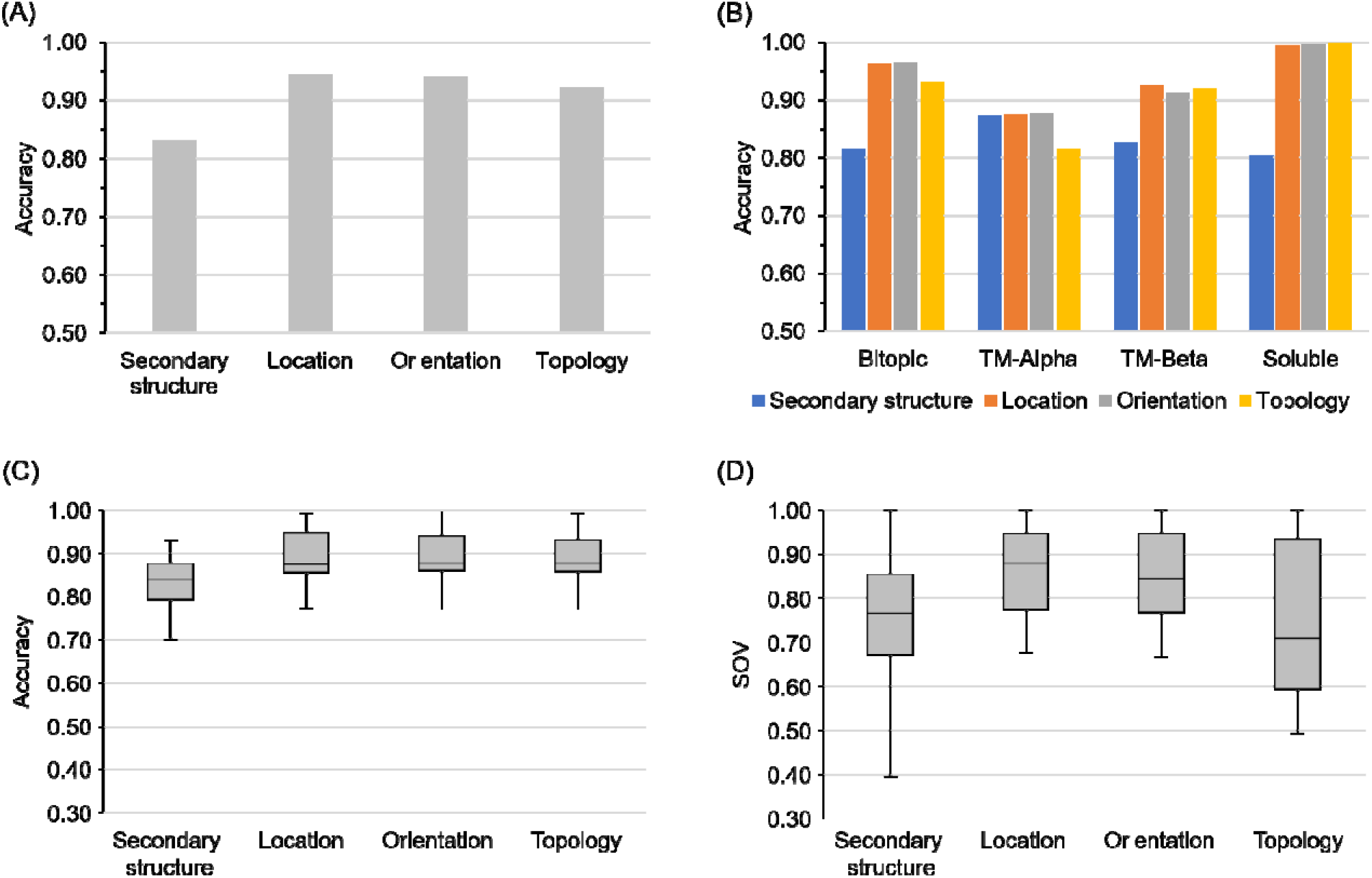
Performance of MASSP on predicting structural features as tested using a blind test set. (A) Overall accuracy achieved by MASSP for each target attribute. (B) Overall accuracy for each target attribute decomposed according to protein class. (C) Distribution of prediction accuracy for each target attribute on test set proteins. (D) Distribution of the segment overlap (SOV) metric for each target attribute on test set proteins.

We also computed the distributions of the Q3 and the SOV measures on test set proteins (Fig. 3C and 3D). For secondary structure prediction, the median Q3 and SOV are 0.830 and 0.766, respectively. For the other three membrane protein-related attributes, the median Q3s are all above 0.875 and the median SOV is 0.879 for location and 0.845 for orientation. However, the median SOV is substantially worse (0.709) for topology prediction. Achieving high SOV for transmembrane topology prediction is difficult because a flipped prediction will result in an essentially zero SOV.

### Near-perfect prediction of protein classes via LSTM

Knowing the structural class of a protein (soluble vs. transmembrane, TM-alpha vs. TM-beta, and bitopic vs. multi-spanning) is essential toward understanding its potential cellular function. MASSP builds on the residue-level predictions of secondary structures, membrane associations, residue orientations, and transmembrane topologies and leverages the power of the LSTM recurrent neural network architecture (Hochreiter and Schmidhuber, 1997) to accurately predict protein classes. In recurrent neural networks, each sequence is treated as a single data point and is processed by iterating through the sequence elements and maintaining a state containing information relative to what it has seen so far. When applied to protein class prediction, the network maintains information about all residues in sequence.

To achieve this, we tokenize protein sequences at residue level by representing each residue with a “word” composed of their residue-level predictions. For instance, a residue predicting to be helix-forming, transmembrane, going toward the cytoplasmic side, and whose sidechain is lipid facing would be represented by the word “HMLD”, and a strand-forming residue in a soluble protein would be represented by the word “ESSS”. We used the training set to both learn a task-specific word-embedding space and train the LSTM model. The hyperparameters of the LSTM model were tuned using the validation set.

The performance the LSTM model was estimated using the test set. As the LSTM model was trained to four protein classes, we report its performance using the Q4 accuracy. Overall, the LSTM model correctly predicted the types of 70 out the 73 proteins in the test set, corresponding to a Q4 accuracy of 95.9% (Table 1). Specifically, the LSTM predicted 36 out of the 37 soluble proteins as soluble (97.3%), 6 out of 8 of the TM-beta proteins to be TM-beta (75%). Remarkably, all TM-alpha proteins are predicted to be TM-alpha, and all bitopic proteins are also predicted to be bitopic.

**Table 1.** Summary of the performance of the LSTM-based protein class predictor.

|  | Bitopic | TM-alpha | TM-beta | Soluble |
| --- | --- | --- | --- | --- |
| Bitopic | 5 | 0 | 0 | 0 |
| TM-alpha | 0 | 23 | 0 | 0 |
| TM-beta | 0 | 0 | 6 | 2 |
| Soluble | 1 | 0 | 0 | 36 |

\* Row headers indicate experimental structural attributes; column headers indicate predicted structural attributes; numbers in the matrices are residue counts.

### MASSP achieves state-of-the-art performance

To evaluate the performance of MASSP relative to other methods in the field, we first compared it to Jufo9D previously developed in our group and three other popular secondary structure prediction methods (PSIPRED, RaptorX-Property, and SPINE-X). Jufo9D is based on a densely connected one-hidden layer feedforward neural network that outputs a 3 × 3 probability matrix for each residue to be in one of three secondary structure types (helix, strand, coil) and one of three transmembrane environment types (membrane core, interface, solution) (Leman et al., 2013). The PSIPRED secondary structure predictor was developed over two decades ago (Jones, 1999) and is still a very popular tool (Buchan and Jones, 2019). PSIPRED consists of two successive densely connected neural networks with input taken in a window of 15 residues and outputs a three-state prediction for the central residue. RaptorX-Property is a web server that employs deep convolutional neural fields to predict secondary structure (SS), solvent accessibility (ACC) and disorder regions (DISO) (Wang et al., 2016a). SPINE-X is a multistep neural-network algorithm that couples secondary structure prediction with prediction of solvent accessibility and backbone torsion angles in an iterative manner (Faraggi et al., 2012). We compared MASSP with PSIPRED, RaptorX-Property and SPINE-X for three-state secondary structure prediction on the test set using the Q3 and the SOV measures.

MASSP achieved the best performance of all secondary structure prediction methods tested in terms of both Q3 and SOV (Figure 4). MASSP achieved a mean Q3 of 0.830, which is comparable to PSIPRED (0.829), but higher than SPINE-X (0.813, p = 0.11, two-tailed paired Student t-test), and RaptorX-Property (0.741, p = 5.8e-11, two-tailed paired Student t-test). Our previous method Jufo9D achieved a moderate median Q3 of 0.762 (p = 2.2e-10, two-tailed paired Student t-test). Again, with a mean SOV of 0.752, MASSP achieved better performance than the four other methods: PSIPRED (0.675, p = 3.5e-5, two-tailed paired Student t-test) and SPINE-X (0.698, p = 2.6e-3, two-tailed paired Student t-test) performed similarly in both evaluations, while RaptorX-Property (0.651, p = 3.4e-7, two-tailed paired Student t-test) was significantly worse. The mean SOV of Jufo9D, which is 0.601 (p = 9.1e-12, two-tailed paired Student t-test), is the lowest among all five methods.

**Figure 4.**
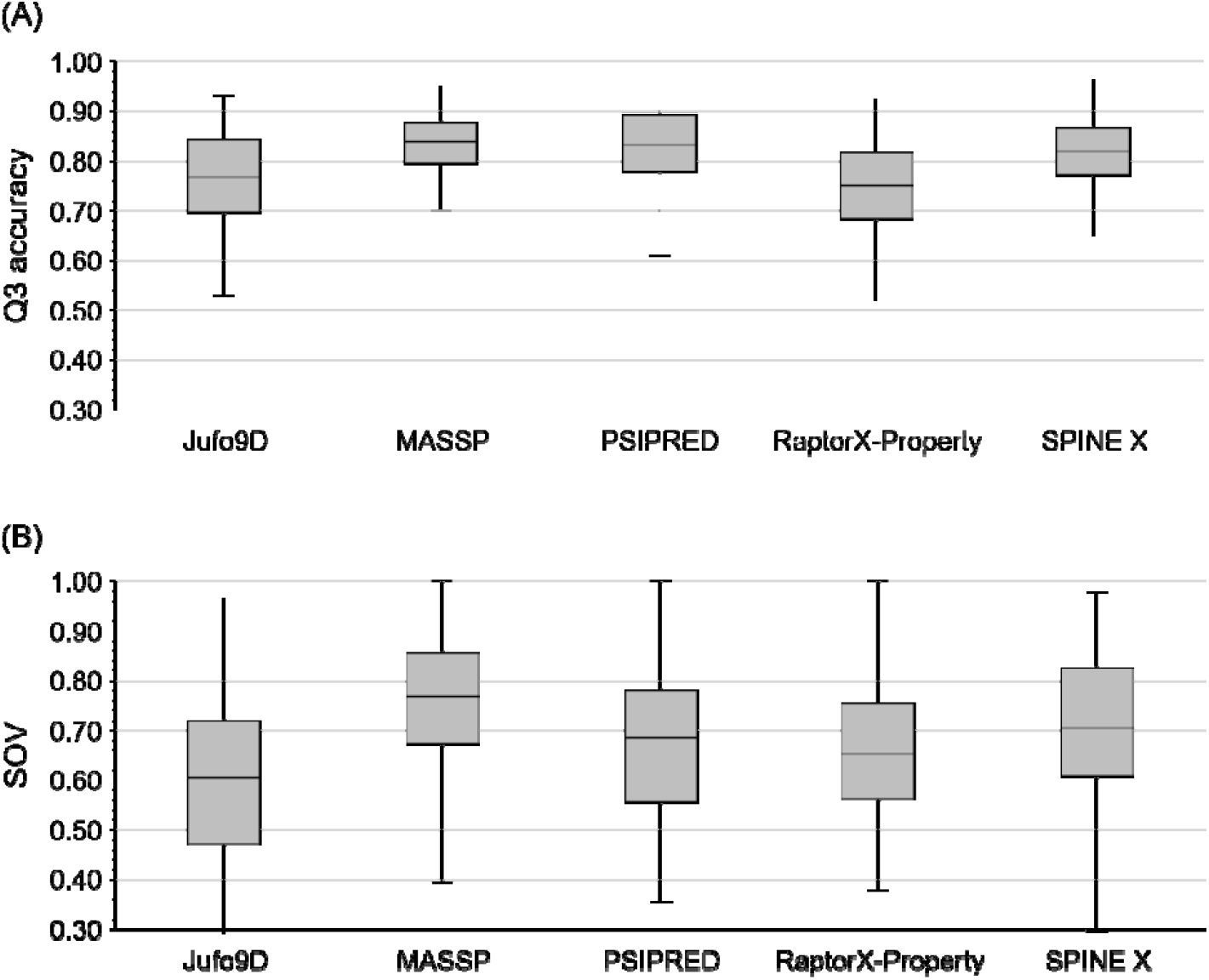
MASSP outperforms Jufo9D, PSIPRED, RaptorX-Property, and SPINE-X at secondary structure prediction. (A) Distribution of Q3 secondary structure accuracies of MASSP on the test set compared to four representative methods. (B) Distribution of performance at secondary structure prediction in terms of the SOV metric for MASSP and four representative methods.

We then compared the performance of MASSP with Jufo9D and other three commonly used methods (MEMSAT3, OCTOPUS, and TMHMM2) at predicting transmembrane segments and topology for TM-alpha proteins. MEMSAT3 is a program which employs a neural network to predict the topology of TM-alpha proteins based on the recognition of topological models and a dynamic programming algorithm to recognize membrane topology models by expectation maximization (Jones, 2007). OCTOPUS uses a combination of artificial neural networks and hidden Markov models to make topology prediction (Viklund and Elofsson, 2008). It first performs a homology search using BLAST to create a sequence profile. This is used as the input to a set of neural networks that predict both the preference for each residue. In the third step, these predictions are used as input to a two-track hidden Markov model, which uses them to calculate the most likely topology (Viklund and Elofsson, 2008).

As shown in Table 2, all five methods achieved a better than 90% accuracy in predicting the number of TM segments for TM-alpha proteins. With a 99.3% accuracy, MASSP achieved the best performance: ∼4-5% higher than MEMSAT3 and ∼8-9% higher than Jufo9D and TMHMM2. Specifically, MASSP correctly predicted 151 TM segments (99.3%), the topology of 23 proteins (82.1%). For topology, MASSP correctly predicted 23 proteins (82.1%) while OCTOPUS predicted 24 (85.7%), MEMSAT3 predicted 23 (82.1%), and TMHMM2 predicted 18 (64.3%). MASSP attained a Q3 of 0.867, and SOV of 0.863. In comparison, OCTOPUS had the highest scores for these metrics with a Q3 of 0.884 and a SOV of 0.867.

**Table 2.** Comparison on the performance of TM segment and topology prediction for TM-alpha proteins.

| Method | Correct #TM | Correct topology | Q3 | SOV |
| --- | --- | --- | --- | --- |
| Jufo9D | 139 (91.4%) | NA | NA | NA |
| MASSP | **151 (99.3%)** | 23 (82.1%) | 0.867 | 0.863 |
| MEMSAT3 | 144 (94.7%) | 23 (82.1%) | 0.871 | 0.839 |
| OCTOPUS | 145 (95.4%) | **24 (85.7%)** | **0.884** | **0.867** |
| TMHMM2 | 137 (90.1%) | 18 (64.3%) | 0.790 | 0.772 |

NA: not available, Jufo9D does not predict residue-level three-state topology.

We next compared the performance of MASSP at predicting transmembrane segments of TM-beta proteins with BOCTOPUS2 (Hayat et al., 2016) and PRED-TMBB2 (Tsirigos et al., 2016), two recently developed methods that were shown to have improved performance over previous methods. As shown in Table 3, MASSP is the best performing method. Both MASSP and BOCTOPUS2 correctly predicted 98 of the 102 transmembrane beta strands and the topology of 6 of the 8 TM-beta proteins in the test set. Whereas PRED-TMBB2 only predicted 88 transmembrane strands correctly and it also inaccurately predicted the topology for one more protein. In terms of Q3 and SOV, the performance of MASSP is substantially better (0.768 and 0.795 respectively) than both BOCTOPUS2 (0.696 and 0.745) and PRED-TMBB2 (0.714 and 0.716).

**Table 3.**
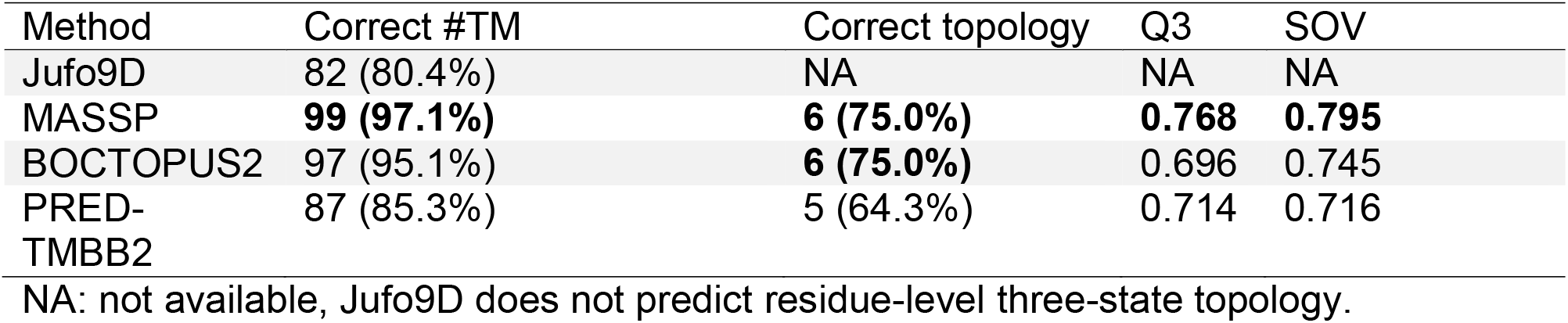
Comparison on the performance of TM segment and topology prediction for TM-beta proteins.

| Method | Correct #TM | Correct topology | Q3 | SOV |
| --- | --- | --- | --- | --- |
| Jufo9D | 82 (80.4%) | NA | NA | NA |
| MASSP | **99 (97.1%)** | **6 (75.0%)** | **0.768** | **0.795** |
| BOCTOPUS2 | 97 (95.1%) | **6 (75.0%)** | 0.696 | 0.745 |
| PRED-TMBB2 | 87 (85.3%) | 5 (64.3%) | 0.714 | 0.716 |

NA: not available, Jufo9D does not predict residue-level three-state topology.

Our extensive comparison on secondary structure and membrane association/topology prediction tasks demonstrates that MASSP performance is comparable to or better than state-of-the-art methods, while being more versatile and application to any protein sequences.

### Examples of accurate MASSP predictions

We demonstrate the predictions made by MASSP on two proteins by mapping secondary structures and topologies predicted by MASSP onto tertiary structures (Fig. 5). We picked the subunit B from the caa3-type cytochrome oxidases from *Thermus thermophilus* (PDB ID: 2yevB) and the TamA protein from the *E. coli* (PDB ID: 4c00A) as two representative examples, because they both consist of large transmembrane and soluble domains and all three types of secondary structures are also well represented. The subunit B from the caa3-type cytochrome oxidases consists of an alpha-helical TM domain and a large extracellular soluble domain, resolved at a resolution of 2.4 Å. TamA is an *E. coli* Omp85 protein involved in autotransporter biogenesis. It comprises a 16-stranded transmembrane β-barrel and three cytoplasmic POTRA domains, resolved at a resolution of 2.3 Å. For 2yevB, MASSP achieved a Q3 of 0.922 and a SOV of 0.908 for secondary structure prediction, and Q3 of 0.987 and a SOV of 0.925 for topology prediction. For 4c00A, MASSP achieved a Q3 of 0.819 and a SOV of 0.843 for secondary structure prediction, and Q3 of 0.940 and a SOV of 0.995 for topology prediction. These two examples illustrate that even for proteins composed of mixed large transmembrane and soluble domains, MASSP can accurately predict both secondary structures and transmembrane topologies.

**Figure 5.**
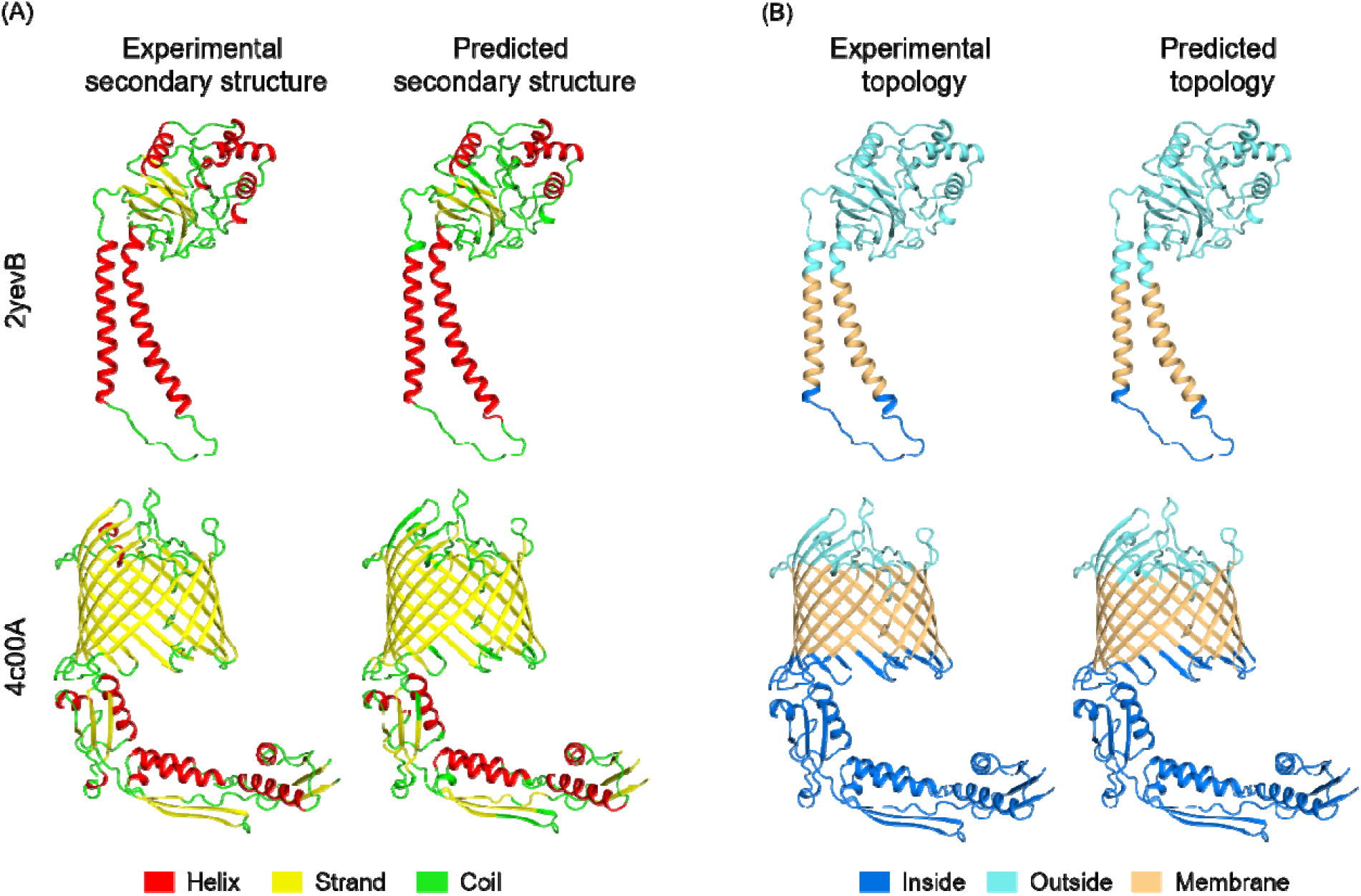
Examples of accurate secondary structure and transmembrane topology prediction by MASSP. (A) Comparisons of experimental and predicted secondary structures mapped to tertiary structures for cytochrome C oxidase subunit 2 (PDB: 2yevB) and translocation and assembly module subunit TamA (PDB: 4c00A) illustrating high accuracy. (B) Comparisons of experimental and predicted residue topologies mapped to tertiary structure for the same proteins.

## Discussion

Prediction of secondary structures and transmembrane regions are fundamental problems in bioinformatics with broad downstream applications. Decades of research into these problems have produced dozens of methods that can be broadly categorized into knowledge-based analyses, generative probabilistic modeling, and discriminative machine learning. However, methods with better performance are still needed giving the increasingly large gap between the set of known sequences and known structures (Koehler Leman et al., 2015; Yang et al., 2018). Among machine-learning methods, those mainly based on neural networks, have been shown to generally perform best, with Q3 accuracies for secondary structure prediction around 80% (Yang et al., 2018); however, their accuracies in predicting transmembrane segments and topologies have been low (Koehler Leman et al., 2015). A newly sequenced protein can be of any one of the structural classes of soluble, bitopic, TM-alpha, or TM-beta. Thus, methods are needed for simultaneous prediction of secondary structures, transmembrane regions, topologies, and protein class.

In this work, we have introduced MASSP, is a multi-task deep-learning framework designed for simultaneous prediction of secondary structures, transmembrane regions, topologies, and protein classes. MASSP simultaneously predicts 1D structural attributes for all four classes of proteins. The core algorithm behind MASSP is a two-tier deep neural network. The first tier is a multi-layer multi-task CNN that predicts residue-level 1D structural attributes. The second tier is a LSTM neural network that treats the predictions of the first tier from the perspective of natural language and predicts the protein class of the input amino acid sequence. We demonstrated that MASSP achieves Q3s of 0.830, 0.942, and 0.922 for predicting secondary structures, residue orientations, and residue topologies, respectively. We also show that the performance of MASSP is comparable or better than several widely used methods for secondary structure and membrane topology prediction.

In addition to performance that is better than or comparable to existing methods, the MASSP framework has other important strengths. First, the multi-tasking nature of MASSP makes it a versatile tool that simultaneously predicts residue-level 1D structural features. The fact that MASSP does not make *a priori* assumptions about whether the given amino acid sequence represents a TM-alpha protein, TM-beta protein, or a soluble protein gives the user a one-stop shop where the secondary structures and membrane associations of an given amino acid sequence of any protein class can be predicted. Thus, MASSP could be applied to the wholly sequenced proteome of an organism to predict the protein class composition of the proteome as well as residue-level secondary structure and membrane association predictions. Second, the multi-output architecture of the CNN would allow us to easily extend MASSP to predict structural properties, such as solvent accessibility and contact order, from amino acid sequence without the requirement of training a different network for each target property.

Despite half-a-century of research into the problems of secondary structure prediction and transmembrane segment and topology prediction, these problems are still open and the field seems to have reached a bottleneck where improvement in prediction accuracy is small and progress is slow (Yang et al., 2018). It strikes us as remarkable that the approach that we took in developing MASSP, i.e. learning patterns in PSSMs with deep CNNs, achieved comparable or better performance than previously developed, more sophisticated methods. This can be partially attributed to applying the “right” type of neural network to the “right” representation of input features, i.e. we leveraged the image-processing power inherent in CNNs and the fact that PSSMs can be thought of images with real-valued “pixels”. Recent advances in protein tertiary structure prediction also demonstrated that a boost in performance can be achieved by developing novel neural network architectures that leverage the problem’s fundamental mathematical essence (AlQuraishi, 2019, 2020; Senior et al., 2020; Xu, 2019). For example, AlphaFold, the algorithm that outperformed all entrants in CASP13, relies on the fundamental principle of the interconvertibility of probability and energy, and predicts the probability distributions of residue pair distances and converts distance distributions to energy landscapes (AlQuraishi, 2020; Senior et al., 2020). These advances in tertiary structure prediction together with our work here suggest that completely solving these problems will likely require the development novel neural network architectures and representation of input features.

## Methods

### Datasets

The sets of multi-spanning α-helical, bitopic, β-barrel, and peripherical proteins were all obtained from the OPM database (Lomize et al., 2006). The datasets were pruned to 25% sequence identity using the PISCES server (Wang and Dunbrack, 2003), and structures for which the resolution is worse than 3.0 Å or were not determined using X-ray crystallography were excluded. The final datasets consist of 241 TM-alpha protein chains, 54 bitopic, and 77 TM-beta protein chains. This dataset was augmented by adding 372 (accounting for 50% of the dataset) soluble protein chains. The set of soluble protein chains was randomly selected from the set of all X-ray only soluble protein chains with resolution better than 3.0 Å and culled at 25% sequence identity using the PISCES server (Wang and Dunbrack, 2003). The dataset was split according to the ratio 8:1:1 as training:validation:test, resulting three subsets containing 165080, 18293, 18571 training examples (residues) for which all four target attributes are available. For alpha-helical membrane proteins, there are 1481, 150, and 158 transmembrane helices in the training, validation, and test sets respectively, and for TM-beta proteins, there are 878, 95, and 104 membrane spanning beta strands respectively. The reference secondary structure elements for each chain were derived from the consensus identification of DSSP (Kabsch and Sander, 1983), Stride (Heinig and Frishman, 2004), and PALSSE (Majumdar et al., 2005). The reference residue location and topology annotations for each chain were derived from the coordinates and membrane boundaries provided by OPM (Lomize et al., 2012).

### Determining beta-strand orientation

The topology of each β-strand composing a barrel was identified by computing the difference in Z-coordinate between the first and last residue in the strand, with strands that ascend in the Z-axis labeled as up (U), while those that descend in Z-axis coordinate labeled as down (D). For orientation, residues were labeled based on whether the C_α_–C_β_ vector points to the pore (P), bilayer (B), or is in solution (S).

Conventionally, three-state Hidden Markov models use interior/exterior/bilayer states and infer strand direction afterwards. We found this state separation to be unappealing for use in training neural networks because in several known structures there are trans-pore segments that cross from intracellular to extra-cellular space, rendering assignment of any residue connected to a pore-crossing segment as intracellular or extracellular ambiguous. Creating an intra-pore state is similarly unappealing due to known structures with dynamic helical plugs that occupy several states. Conversely, we are aware of no beta barrels in which the barrel flips under physiologically relevant conditions, leaving the assignment of U/D/S states unambiguous.

### PSSM calculation

The position-specific-scoring matrix (PSSM) of an amino acid sequence contains the log-odds of each of the 20 amino acids observed at each of the sequence positions over evolutionary timescale (Altschul et al., 1997). To compute the PSSM for a target amino acid sequence, we first queried the UniRef20 protein sequence database for sequences homologous to the target sequence using the HHblits method (Remmert et al., 2012). The parameters used in the searching were three iterations with a e-value of 0.001, a maximum sequence identity of 0.90, a minimum coverage of 0.50 of the target sequence residues. The multiple sequence alignment generated by running HHblits was used as input to an in-house C++ utility software implementing the algorithm described in (Altschul et al., 1997) to compute the floating point-valued PSSM for the target sequence.

### Network architecture

The performance of a neural network depends on many hyperparameters specifying the architecture of the neural network. In designing MASSP, we considered the number of convolutional and densely connected layers, the number of units in each layer, and the size of the input feature matrix. We tuned hyperparameters by training models using the training set and comparing model performances on the validation set. We note that the test set was never touched before the final model was selected.

We followed a workflow of developing deep-learning models recommended in (Chollet, 2018). Specifically, we started out with a very small architecture that has a total of 1339 trainable parameters. This model takes a 7 × 20 input feature matrix, one convolutional layer with weight ReLU units, and one densely connected layer with eight ReLU units. Our goal at this stage was to have a model that is better than a baseline model, i.e. to make sure that patterns in the input PSSM can be learned by the chosen type of neural network. In fact, after 20 epochs of training, we observed that the validation loss was still decreasing (Fig. S1A), indicating that a model with a larger architecture may achieve higher performance.

We next scaled up the architecture of the model by adding two additional convolutional layers, one additional layer of densely connected units, and increasing the number of ReLU units in each layer to 16, 32, 64, 32, and 64, respectively. We also expanded the size of the input feature matrix to cover 15 residues on each side of the central residue. The goal was to have a model that is sufficiently powerful. This was confirmed by observing the model’s performance on the validation set began to degrade only after the first four epochs of training (Fig. S1B).

Starting with this sufficiently powered model, we continued several rounds of hyperparameter tuning by scaling down the size of the architecture and adding regularization (see Fig. S1C and S1D for two examples). In the end, we selected a model with the flattened layer regularized by a 0.25 dropout rate and that appeared to have the optimal overall prediction accuracy on the validation set. The architecture of the final model is shown in Fig. 1B.

### Network construction and training

MASSP is a two-tier neural network system designed for simultaneous prediction of four 1D structural attributes and the protein class for the input amino acid sequence. The first tier is a multi-output deep CNN and was constructed using the functional API of the Keras model-level deep-learning framework for Python, with TensorFlow 2.0.0 serving as the backend engine. The output layer of this multi-output CNN consists of four heads each representing a separate one of the four target structural attributes. The final model was trained using the Adam optimizer (Kingma and Ba, 2014) for 10 epochs with default hyperparameters (maximum learning rate = 0.001, *β*_l_ = 0.9, *β*_2_ = 0.999. Kernel weights of the model were initialized using the Glorot uniform initializer and updated after processing each batch of 64 training examples. The activation of the last layer was the softmax function and the loss function was the categorical cross entropy function for all output heads. By default, Keras sums all four losses into a global loss that is back propagated to update trainable weights of the CNN. We did not reweight individual losses as no one target attribute can be said important than another.

### Generating predictions using other methods

PSIPRED and MEMSAT3 predictions were generated by running PSIPRED and MEMSAT3 on test set protein sequences locally with the recommended UniRef90 sequence database (PSIPRED version 4.02, MEMSAT version 3.0). SPINE-X predictions were similarly generated by running SPINE-X 2.0 locally with the UniRef90 sequence database. RaptorX-Property predictions were generated by running RaptorX-Property version 1.02 on test protein sequences with the recommended UniRef20 sequence database. Source code of these locally run methods were downloaded and compiled locally, and the programs were set up following recommendations. TMHMM2 predictions were generated by submitting test sequences to the TMHMM web server at http://www.cbs.dtu.dk/services/TMHMM/. Similarly, OCTOPUS predictions were obtained from its web server at http://boctopus.bioinfo.se/octopus/. BOCTOPUS2 predictions were generated by submitted all test protein sequences to the BOCTOPUS2 web server at http://boctopus.bioinfo.se/pred/ with default settings. PRED-TMBB predictions were generated by submitting all test protein sequences to the PRED-TMBB web server at http://www.compgen.org/tools/PRED-TMBB2 in single-sequence mode with default settings.

### Performance measures

We employed two of the most commonly used performance measures to evaluate MASSP and to compare it with other methods. The first measure, called the Q3 accuracy, is defined as the fraction of residues for which the three-state target attributes are correctly predicted. For secondary structure prediction in this work,

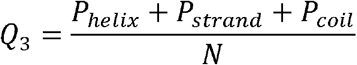

where *N* is the total number of predicted residues and *P*_*x*_ is the number of correctly predicted secondary structures of type *x*. The accuracy measure for evaluating the prediction of location, orientation, and membrane topology were similarly defined.

The other measure is known as the fractional overlap of segments (SOV) originally proposed by (Rost et al., 1994). SOV measures the percentage of correctly predicted secondary structure segments rather than individual residue positions, and it pays less attention to small errors in the ends of structural elements. In this work, we used an updated definition of SOV introduced by (Zemla et al., 1999) and implemented in Perl by (Liu and Wang, 2018). For secondary structure prediction,

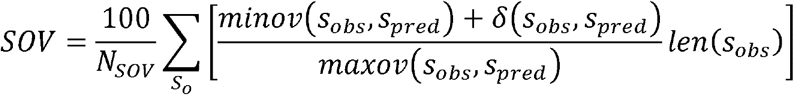

where *s*_*obs*_and *s*_*pred*_represent all observed and predicted segments of helices, strands, and coils. is the set of all overlapping pairs of *s*_*obs*_ and *s*_*pred*_where the segments are the same state. is the length in residues of any segment *s*_*obs*_; *minov*(*s*_*obs*,_ *s*_*pred*_) is the length of the actual overlap between any segment pair (*s*_*obs*_, *s*_*pred*_) in *S*_*0*_and *maxov*(*s*_*obs*_, *s*_*pred*_) is the total extent to which at least one residue is that state. *N*_*sov*_ is the total number of residues in *s*_*obs*_ in all pairs in plus the number of residues in any *s*_*obs*_ that are not overlapped by a predicted segment of the same state. The summation represents the fraction of the segment pair that the observed and predicted states agree. *δ*(*s*_*obs*_, *s*_*pred*_) is added to allow for some variation in segment boundaries and is defined as

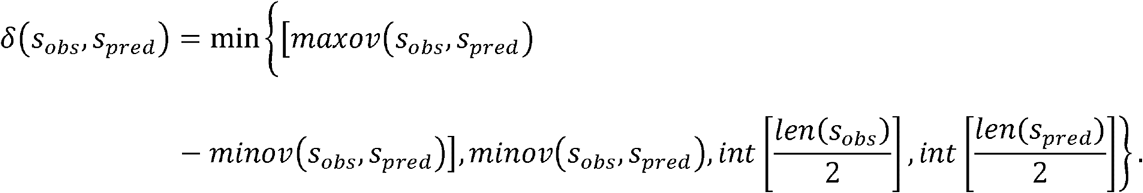

While SOV is typically used to evaluate performance on secondary structure prediction, it is a general measure that is well suited for, and thus was also used in, evaluating prediction of the other three structural attributes in this study.

### Availability

MASSP is publicly available at http://www.meilerlab.org/index.php/servers/show?s_id=26. In addition to this web server, all raw data and code are also available as supplementary data to this manuscript. The C++ utility for computing position-specific-scoring matrices using multiple sequence alignments was implemented in the sequence analysis module of the BioChemical Library software, source and executables for which are available free of charge for academic use from www.meilerlab.org/bcl_academic_license.

## Supporting information

Code and Raw Data

## Author contribution

B.L., J. Mendenhall, and J. Meiler conceived the study. B.L. and J. Mendenhall developed the method and performed data analysis. J. Meiler supervised the project. All authors read, commented, and edited the manuscript.

## Acknowledgement

This work was supported by NIH awards R01 GM080403, R01 HL122010, R01 DA046138, and by the Deutsche Forschungsgemeinschaft (DFG, German Research Foundation) through SFB1423, project number 421152132, subproject Z04, A07. This work was also conducted in part using the computational resources of the Advanced Computing Center for Research and Education at Vanderbilt University.

