## Supporting Information for "A multi-task deep-learning system for predicting membrane associations and secondary structures of proteins"

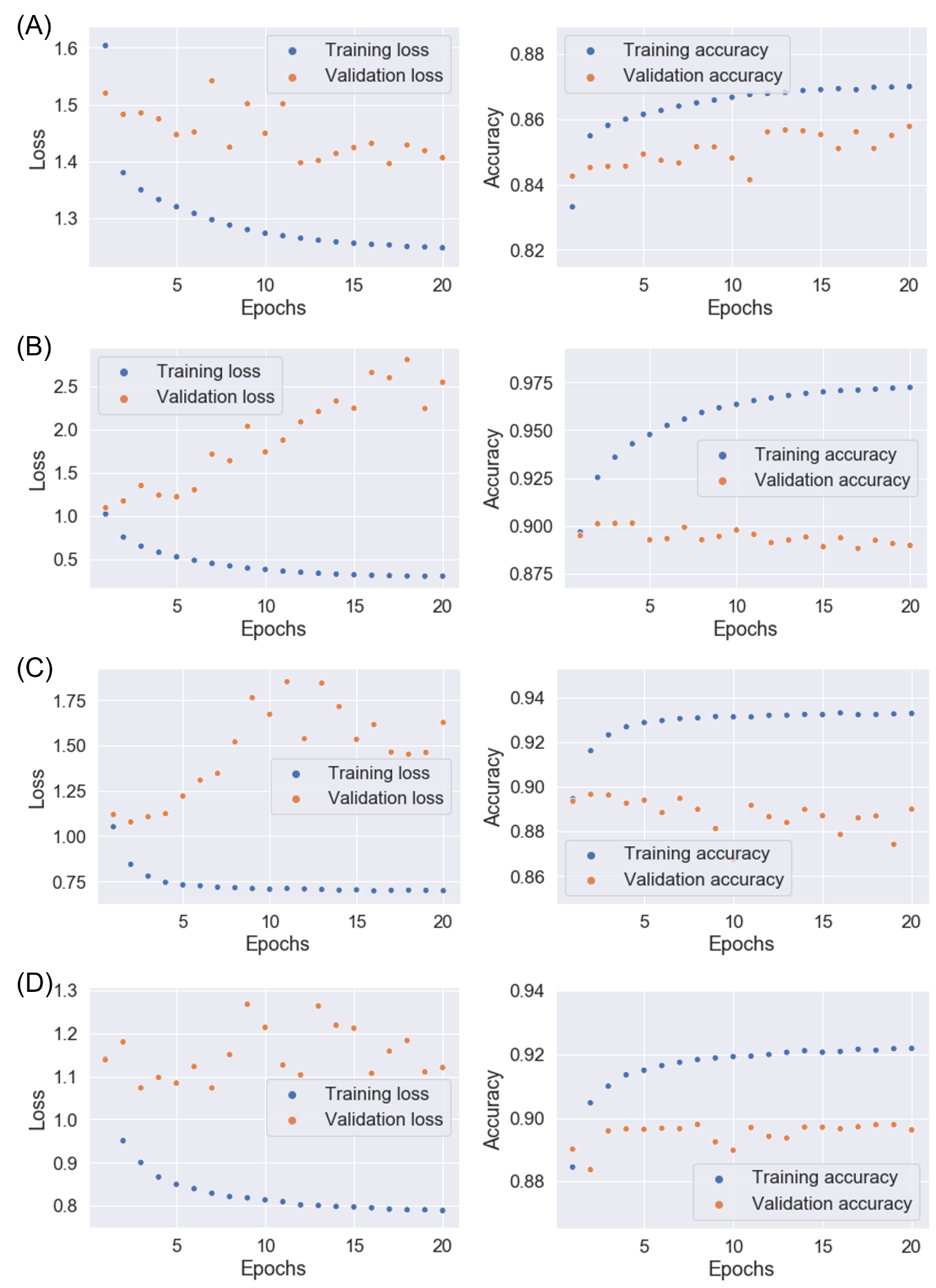

**Figure S1. Tuning architectural hyperparameters of the CNN component of MASSP.** (A) The training and validation losses and accuracies of a starting “small” model with 1339 trainable parameters. This model takes a $7\times21$ input matrix and consists of one convolutional layer and one dense layer, both with eight ReLU units. This model was underfitting because validation loss continued to drop, and validation accuracy continued to increase. (B) The training and validation losses and accuracies of a sufficiently powerful, overfitting model with 198187 trainable parameters. This model takes a $31\times21$ input matrix and consists of three convolutional layers and two densely connected layers with 16, 32, 64 32, and 64 ReLU units, respectively. It is immediately obvious that this model was overfitting. (C). The same architecture as in (B) but a dropout layer with 0.25 rate was added before each dense layer. The model was still overfitting after adding dropout regularization, indicating that the architecture was too large. (D) The training and validation losses and accuracies of a model with 34755 trainable parameters. This model takes a $15\times21$ input matrix and consists of two convolutional layers and one densely connected layer with 16, 24, and 32 ReLU units, respectively. Overfitting was more or less remedied in this model. We continued several more rounds of hyperparameter tuning and selected a model shown in Fig. 1B that appeared to have optimal performance on the validation set.

**Table S1. Five-letter PDB IDs of training set protein subunits grouped according to protein class.**

| Bitopic | | | | | | | | |
| --- | --- | --- | --- | --- | --- | --- | --- | --- |
| 1ezvH | 1oqwA | 2dyrK | 2j7aI | 3cx5H | 3wfdC | 3wu2X | 4pj0R | 5l8rH |
| 1jb0F | 1pp9G | 2dyrL | 2j8cH | 3cx5I | 3wu2E | 4kytB | 4pj0Y | 5mrwC |
| 1jb0J | 2dyrD | 2dyrM | 2xfnA | 3jqoA | 3wu2F | 4ogqC | 4wmzA | 5y5s0 |
| 1m56D | 2dyrI | 2fyuK | 3cx5D | 3s8gB | 3wu2H | 4ogqD | 5awwE | 6mjpC |
| 1nkzA | 2dyrJ | 2j58A | 3cx5E | 3vmtA | 3wu2J | 4pj0K | 5hk1A |  |
| TM-alpha | | | | | | | | |
| 3kcuA | 3ug9A | 4ikvA | 4rdqA | 5a1sC | 5jjeB | 5uldA | 6bmmA | 6i1rA |
| 3ldcA | 3v5uA | 4jr9A | 4ri2A | 5a43A | 5jnqA | 5uniA | 6bw6A | 6i6hA |
| 3llqA | 3w4tA | 4k1cA | 4rp9A | 5ajiA | 5jsiA | 5uniB | 6cb2A | 6i9kA |
| 3m73A | 3wajA | 4m48A | 4ryoA | 5awwG | 5kbwA | 5v7pA | 6d0jA | 6idpA |
| 3mp7A | 3wfdB | 4m5bA | 4tq4A | 5awwY | 5kukA | 5v8kA | 6d91A | 6igkA |
| 3n5kA | 3wu2A | 4mndA | 4twkA | 5ax0A | 5lwyA | 5vrhA | 6e9nA | 6iu3A |
| 3ne5A | 3wu2B | 4n6hA | 4u9nA | 5aynA | 5mlzA | 5wudA | 6ebuA | 6iyxA |
| 3puwF | 3wu2D | 4n7wA | 4uc1A | 5azbA | 5mrwA | 5x3xQ | 6edqA | 6m96B |
| 3puwG | 3wu2Z | 4o6mA | 4umwA | 5bz3A | 5mrwB | 5xapA | 6ei3A | 6mjpF |
| 3rkoA | 4a01A | 4o6yA | 4v1fA | 5c6pA | 5nv9A | 5xj5A | 6eyuA | 6mjpG |
| 3rkoJ | 4bemJ | 4ogqB | 4wd8A | 5c78A | 5o0tA | 5xlsA | 6f2gA | 6mwaB |
| 3rkoK | 4cadC | 4p02A | 4x5mA | 5dirA | 5oc0A | 5y50A | 6ffvA | 6nc9A |
| 3rkoL | 4d2eA | 4p79A | 4xesA | 5dqqA | 5oglA | 5y79A | 6fl9A | 6o3cA |
| 3rkoM | 4dveA | 4pgrA | 4xk8G | 5dwyA | 5oqtA | 5ys3A | 6fmxA | 6ob7A |
| 3rkoN | 4dx5A | 4phzA | 4xtlA | 5edlA | 5sv0A | 6a2jA | 6fv8A | 6oh3A |
| 3rlbA | 4eiyA | 4phzB | 4xu4A | 5ezmA | 5svkA | 6a2wA | 6g94A | 6or2A |
| 3rqwA | 4g7vS | 4phzC | 4yl3A | 5fgnA | 5sytA | 6a6mA | 6g9xA | 6rnkA |
| 3s8gA | 4gx0A | 4pl0A | 4ymkA | 5gufA | 5tcxA | 6ak3A | 6h2fA |  |
| 3tdsA | 4hfiA | 4qndA | 4ymuC | 5i20E | 5tgzA | 6al2A | 6h59A |  |
| 3tijA | 4huqS | 4qtnA | 4zp0A | 5iwsA | 5tinA | 6b87A | 6h7dA |  |
| 3tuiA | 4huqT | 4quvA | 4zr1A | 5j4iA | 5tsaA | 6barA | 6hd8B |  |
| 3tx3A | 4i0uA | 4r0cA | 4zwcA | 5jeqA | 5uiwA | 6bd4A | 6hlpA |  |
| TM-beta | | | | | | | | |
| 1ek9A | 1yc9A | 2x27X | 3dwoX | 3o44A | 4afkA | 4rjwA | 5fvnA | 6fokA |
| 1fepA | 2ervA | 2x55A | 3dzmA | 3pguA | 4e1sA | 4rl8A | 5ldvA | 6gieA |
| 1kmoA | 2gr7A | 2x9kA | 3emnX | 3qq2A | 4fuvA | 4rlcA | 5mdrA | 6hcpA |
| 1p4tA | 2gufA | 2ynkA | 3fhhA | 3rfzB | 4geyA | 4y25A | 5o65A | 6i96A |
| 1qd6C | 2porA | 3aehA | 3fidA | 3v8xA | 4n75A | 5dl7A | 6e4vA | 6qgwA |
| 1qjpA | 2vdfA | 3bs0A | 3gp6A | 3w9tA | 4ql0A | 5dl8A | 6ehbA |  |
| 1uynX | 2wjrA | 3cslA | 3kvnA | 3wi5A | 4rdrA | 5fokA | 6eheA |  |
| Soluble | | | | | | | | |
| 1ayoA | 1t6uA | 2owlA | 3djeA | 3m4aA | 4c4aA | 4neoA | 5d7wA | 6bsqA |
| 1b8kA | 1t82A | 2p06A | 3dsbA | 3mcrA | 4cd5A | 4o9dA | 5e7tB | 6dfdA |
| 1c1yB | 1th7A | 2pspA | 3e0rA | 3mtwA | 4cdpA | 4obmA | 5fcmA | 6dnqE |
| 1c3cA | 1unnC | 2q2fA | 3e6qA | 3n0aA | 4cjdA | 4ojdH | 5h66C | 6ea3A |
| 1chmA | 1ut7A | 2q3sA | 3eo7A | 3n0uA | 4cw4A | 4oo3A | 5il5A | 6eerA |
| 1cy5A | 1vkiA | 2q43A | 3eshA | 3n17A | 4dooA | 4pavA | 5jazA | 6es9A |
| 1d2nA | 1x9iA | 2q66A | 3eusA | 3nk6A | 4ebrA | 4pcqA | 5jbxA | 6eyxA |
| 1eejA | 1xnfA | 2qkpA | 3f52A | 3nqiA | 4eqaC | 4pj3A | 5jiwA | 6go4A |
| 1eq2A | 1y66A | 2qmiA | 3fcmA | 3on5A | 4ercA | 4qb7A | 5kc1A | 6gyeA |
| 1fpoA | 1yw4A | 2qs9A | 3fcnA | 3p51A | 4eswA | 4qozC | 5m48A | 6gz8A |
| 1g8pA | 1z0xA | 2qwwA | 3fg8A | 3phsA | 4euoA | 4qq0A | 5m5eB | 6hazA |
| 1h16A | 1za7A | 2r01A | 3gpvA | 3pnrB | 4exoA | 4rbrA | 5mu5A | 6hx0A |
| 1h5wA | 2a1rA | 2r31A | 3gwqA | 3pu9A | 4f3vA | 4rctA | 5n5fA | 6i03A |
| 1hk8A | 2a5yB | 2va0A | 3h7oA | 3qdpA | 4gimA | 4rjjA | 5n7eB | 6ibeA |
| 1hxrA | 2aegA | 2vwsA | 3hbcA | 3qwnA | 4go6B | 4rz4A | 5naaA | 6j0pA |
| 1hypA | 2c0cA | 2wm3A | 3hmsA | 3qwuA | 4h0aA | 4s1pA | 5ncbA | 6km7A |
| 1i7wB | 2c3gA | 2wtmA | 3ho6A | 3qxcA | 4h3vA | 4trtA | 5ngjA | 6n2cA |
| 1im3D | 2cdcA | 2x5yA | 3hoiA | 3r0aA | 4hi8A | 4u4cB | 5o0jA | 6n63A |
| 1j0pA | 2ci1A | 2ybyA | 3hryA | 3ry3A | 4hpmB | 4uabA | 5o7hF | 6nzmA |
| 1jhsA | 2ckkA | 2z3xA | 3ic3A | 3ufbA | 4hrvA | 4v1tA | 5od4A | 6o60B |
| 1k8wA | 2dtjA | 2z84A | 3id6A | 3v7iA | 4htpA | 4w66A | 5qhhA | 6o87A |
| 1ka8A | 2dwuA | 2zxxC | 3ijmA | 3vkwA | 4hvkA | 4w9zA | 5t2pA | 6qj2B |
| 1lniA | 2e11A | 3a1gA | 3ikbA | 3vr0A | 4i3yA | 4waiA | 5th5A | 6qlvA |
| 1n71A | 2efkA | 3ag7A | 3iv6A | 3wqoA | 4ivnA | 4wbeA | 5tt5A | 6rcqA |
| 1nh2C | 2fiyA | 3al2A | 3jqhA | 3wxfA | 4jejA | 4ww7A | 5u96A | 6s07A |
| 1o7dC | 2gnxA | 3aq2A | 3jxoA | 3zpjA | 4kdwA | 4xr9A | 5ufyA | 6uq8A |
| 1p27B | 2hboA | 3bedA | 3jz0A | 3zsjA | 4l0jA | 4xwhA | 5um2A |  |
| 1q74A | 2hcfA | 3bwzA | 3k2oA | 4a15A | 4l0rA | 4y9iA | 5vbnA |  |
| 1r7lA | 2i39A | 3c75H | 3kzqA | 4a6qA | 4l8jA | 4yarA | 5wddA |  |
| 1s5dA | 2i3dA | 3chjA | 3ledA | 4aeqA | 4lfuA | 4ynxA | 5z7bA |  |
| 1sh8A | 2ibdA | 3cj1A | 3lkbA | 4at7B | 4m1uA | 5ajsA | 5zhoA |  |
| 1skvA | 2je6A | 3cjdA | 3lp5A | 4ay0A | 4m8aA | 5azxA | 6a0nA |  |
| 1sqhA | 2o3bB | 3d1cA | 3lufA | 4bwnA | 4mfiA | 5c5zA | 6ae9A |  |
| 1sw6A | 2omzB | 3d3bJ | 3ly7A | 4bwrA | 4n8nA | 5cs0A | 6ahoA |  |

**Table S2. Five-letter PDB IDs of validation set protein subunits grouped according to protein class.**

| Bitopic | | | | | | | | |
| --- | --- | --- | --- | --- | --- | --- | --- | --- |
| 2zxeG | 3mp7B | 4p02B | 5uhqB | 5y5s1 |  |  |  |  |
| TM-alpha | | | | | | | | |
| 1jb0A | 1kf6C | 1m0lA | 1okcA | 1p49A | 1u7gA | 2bl2A | 2dyrA |  |
| 1jb0K | 1kf6D | 1nekC | 1orsC | 1q16C | 2a65A | 2bs2C | 2dyrB |  |
| 1jb0L | 1kqfC | 1nekD | 1otsA | 1u19A | 2bhwA | 2cfqA | 2dyrC |  |
| TM-beta | | | | | | | | |
| 1uunA | 2mprA | 3pikA | 4d5bA | 4fqeA | 5azpA | 5dl5A | 5fq6D |  |
| Soluble | | | | | | | | |
| 1dm9A | 1ydyA | 2ghvC | 3a0oA | 3ushA | 4hjiA | 4ponA | 5munA | 6cb7A |
| 1nh2D | 1yudA | 2obpA | 3kewA | 3vorA | 4i3mA | 4r81A | 5o63A | 6e1zA |
| 1y1xA | 1z70X | 2pd1A | 3oioA | 4bt7A | 4l9aA | 4x5pA | 5w5cE | 6gsoA |
| 1y8aA | 1zpsA | 2z69A | 3pjpA | 4fo9A | 4lzxB | 5mecA | 5x6bE | 6n1bA |
| 7odcA |  |  |  |  |  |  |  |  |

**Table S3. Five-letter PDB IDs of test set protein subunits grouped according to protein class.**

| Bitopic | | | | | | | | |
| --- | --- | --- | --- | --- | --- | --- | --- | --- |
| 1kqfB | 2dyrG | 2zxeB | 5oy0I | 1kqfB |  |  |  |  |
| TM-alpha | | | | | | | | |
| 2h88C | 2jlnA | 2uuhA | 2wswA | 2yevC | 3c02A | 3dh4A | 3h90A |  |
| 2h88D | 2nq2A | 2vpzC | 2xtvA | 2z73A | 3cx5C | 3fb5A | 3k3fA |  |
| 2j8cM | 2qtsA | 2w2eA | 2yevB | 3b9yA | 3d31C | 3giaA |  |  |
| TM-beta | | | | | | | | |
| 3b07B | 3qraA | 4b7oA | 4c00A | 4meeA | 4q35A | 6eusA | 7ahlA |  |
| Soluble | | | | | | | | |
| 1fviA | 2e7jA | 2vqeC | 3ct8A | 3ts9A | 4gudA | 4p8iA | 5j1hA | 5w1uA |
| 1jk0B | 2fhpA | 2xocA | 3fppA | 3vwcA | 4mb0A | 4y97B | 5lsfA | 6cbrA |
| 1k4iA | 2i51A | 3a2zA | 3gwrA | 3zrxA | 4nb5A | 5d23A | 5ugrA | 6detA |
| 1qqfA | 2ra8A | 3ca8A | 3iusA | 3zzsA | 4o06A | 5gofA | 5ugzA | 6evuA |
| 6i4eA |  |  |  |  |  |  |  |  |

**Table S4. Data set summary statistics by residue counts.**

|  | Training | Validation | Test |
| --- | --- | --- | --- |
| Helix | 78415 | 7912 | 8641 |
| Strand | 25923 | 3115 | 3001 |
| Coil | 60742 | 7266 | 7521 |
| Membrane | 42724 | 4640 | 4894 |
| Soluble | 122356 | 13653 | 14269 |
| Up | 20192 | 2238 | 2270 |
| Down | 19986 | 2124 | 2249 |
| Outside | 124902 | 13931 | 14644 |
| TM helix | 1481 | 150 | 152 |
| TM strand | 878 | 94 | 102 |

**Table S5. Q3 values by protein class.**

|  | Bitopic | TM-alpha | TM-beta | Soluble |
| --- | --- | --- | --- | --- |
| Secondary structure | 0.816 | 0.874 | 0.828 | 0.805 |
| Location | 0.964 | 0.876 | 0.927 | 0.996 |
| Orientation | 0.966 | 0.877 | 0.914 | 0.997 |
| Topology | 0.932 | 0.817 | 0.920 | 0.998 |

**Table S6. Contingency matrices for test set predictions.**

|  | Helix | | Strand | | Coil |
| --- | --- | --- | --- | --- | --- |
| Helix | 7278 | | 37 | | 921 |
| Strand | 12 | | 2274 | | 569 |
| Coil | 887 | | 690 | | 5903 |
|  | | **Inside membrane** | | **Outside membrane** | |
| Inside membrane | | 4083 | | 620 | |
| Outside membrane | | 426 | | 13442 | |
|  | **Lipid** | | **Pore** | | **Out** |
| Lipid | 3608 | | 15 | | 524 |
| Pore | 29 | | 399 | | 50 |
| Out | 209 | | 215 | | 13906 |
|  | **Up** | | **Down** | | **Out** |
| Up | 1653 | | 286 | | 274 |
| Down | 234 | | 1565 | | 229 |
| Out | 209 | | 215 | | 13906 |

* Row headers indicate experimental structural attributes; column headers indicate predicted structural attributes; numbers in the matrices are residue counts.
