## Supplementary material for "A multi-task deep-learning system for predicting membrane associations and secondary structures of proteins": Code and Raw Data: convert_massp_to_top.py

```
import sys


top_file = sys.argv[1]
with open(top_file, 'rt') as ipf:
    header = next(ipf).strip()
    top_str = ''.join([l.strip() for l in ipf])

# get first membrane residue direction
for l in top_str:
    if l == 'O':
        continue
    else:
        first_mem_res = l
        break

prev_ltr = ''
new_str = ''
for i, l in enumerate(top_str):
    if l == 'O':
        if prev_ltr == '':
            if first_mem_res == 'D':
                new_str += 'o'
            else:
                new_str += 'i'
        elif prev_ltr == 'D':
            new_str += 'i'
        else:
            new_str += 'o'
    else:
        new_str += 'M'
        prev_ltr = l

with open(sys.argv[2] + '_new_top.fasta', 'wt') as opf:
    opf.write(header)
    opf.write('\n')
    opf.write(new_str)
    opf.write('\n')
```
