## Supplementary material for "A multi-task deep-learning system for predicting membrane associations and secondary structures of proteins": Code and Raw Data: count_correct_tmb.py

```
import sys


true_file = sys.argv[1]
pred_file = sys.argv[2]

with open(true_file, 'rt') as ipf:
    true_str = ''.join([l.strip() for l in ipf if not l.startswith('>')])

# determine TM segment boundaries
tmh_boundaries = []
prev_res = ''
start = 0
end = 0
for i, x in enumerate(true_str):
    if x == 'M' and prev_res != 'M':
        start = i
    if (x != 'M' and prev_res == 'M') or (x == 'M' and i == len(true_str) - 1):
        end = i
    if end != 0:
        tmh_boundaries.append((start, end))
        start = 0
        end = 0
    prev_res = x

with open(pred_file, 'rt') as ipf:
    pred_str = ''.join([l.strip() for l in ipf if not l.startswith('>')])

true_count = 0
pred_count = 0
for start, end in tmh_boundaries:
    if end - start >= 3:
        true_count += 1
    if pred_str[start:end].count('M') >= 3:
        pred_count += 1

print(true_count, pred_count)
```
