## Supplementary material for "A multi-task deep-learning system for predicting membrane associations and secondary structures of proteins": Code and Raw Data: count_correct_tmh.py

```
import sys


true_file = sys.argv[1]
jufo9d_file = sys.argv[2]

with open(true_file, 'rt') as ipf:
    true_str = ''.join([l.strip() for l in ipf if not l.startswith('>')])

# determine TM segment boundaries
tmh_boundaries = []
prev_res = ''
start = 0
end = 0
for i, x in enumerate(true_str):
    if x == 'M' and prev_res != 'M':
        start = i
    if (x != 'M' and prev_res == 'M') or (x == 'M' and i == len(true_str) - 1):
        end = i
    if end != 0:
        tmh_boundaries.append((start, end))
        start = 0
        end = 0
    prev_res = x

with open(jufo9d_file, 'rt') as ipf:
    jufo9d_str = ''.join([l.strip() for l in ipf if not l.startswith('>')])

true_count = 0
jufo9d_count = 0
for start, end in tmh_boundaries:
    if end - start >= 10:
        true_count += 1
    if jufo9d_str[start:end].count('M') >= 10:
        jufo9d_count += 1

print(true_count, jufo9d_count)
```
