## Supplementary material for "A multi-task deep-learning system for predicting membrane associations and secondary structures of proteins": Code and Raw Data: massp.py

```
# -*- coding: utf-8 -*-
"""
MASSP is a two-tier multitasking deep neural network for secondary structure
and transmembrane topology prediction. It makes no a priori assumption about
what protein class the input sequence belongs to. Thus, it is applicable to 
the whole proteome of any organism. Our benchmarking analysis indicates that
MASSP has superior performance to some of the existing best methods: PSIPRED,
RaptorX-Property, SPINE-X, OCTOPUS, MEMSAT3, TMHMM2, BOCTOPUS, PRED-TMBB2.
We're currently preparing a manuscript on this method.

@author: Bian Li
@
"""

import numpy as np
import os, sys
from tensorflow.keras import models
from argparse import ArgumentParser
from tensorflow.keras.preprocessing.sequence import pad_sequences
from tensorflow.keras.preprocessing.text import Tokenizer


# global variables
SEQ_DB = 'path to uniref20 sequence database'
HHBLITS = 'path to hhblits binary'
REFORMAT = 'path to hhblits reformat.pl script'
BCL = 'path to the BioChemical Library (BCL) binary'
PSSM_BIN = 'path to the compute_pssms.sh script'


def run_hhblits(fasta_file, prefix, ncpu='1'):
    """
    Searches sequences that are homologous to the query sequence using
    the HHblits software suite. 

    Parameters
    ----------
    fasta_file : TYPE
        DESCRIPTION.
    prefix : TYPE
        DESCRIPTION.
    ncpu : TYPE, optional
        DESCRIPTION. The default is '1'.

    Returns
    -------
    None.

    """
    hhblits_options = [
        '-i', fasta_file,
        '-d', SEQ_DB,
        '-e', '0.001',
        '-n', '2',
        '-cov', '50',
        '-id', '90',
        '-diff', 'inf',
        'realign_max', '500000',
        '-pre_evalue_thresh', '0.01',
        '-oa3m', prefix + '.a3m',
        '-o', prefix + '.hhr',
        '-ohhm', prefix + '.hhm',
        '-cpu', ncpu
    ]
    
    hhblits_cmd = ' '.join(['hhblits'] + hhblits_options)
    os.system(hhblits_cmd)
    
    # reformatting multiple sequence alignment
    reformat_cmd = ' '.join(
        [
            REFORMAT, '-r',
            'a3m', 'clu',
            prefix + '.a3m',
            prefix + '.clu'
        ]
    )
    os.system(reformat_cmd)
    
    # generate a position-specific scoring matrix
    pssm_cmd = ' '.join(
        [
            BCL, 'bioinfo:MSA2PSSM',
            prefix + '.clu',
            prefix + '.pssm',
            '-efractional_pseudocount', '1',
            '-pseudocount', '1',
            '-msa_background',
            '-fractional_pseudocount', '0.15'
        ]
    )
    os.system(pssm_cmd)


def read_pssm(pssm_file):
    """
    Read a given position-specific scoring matrix into 2D NumPy Array.

    Parameters
    ----------
    pssm_file : TYPE
        DESCRIPTION.

    Returns
    -------
    pssm_rows : TYPE
        DESCRIPTION.
        
    """
    pssm_rows = []
    with open(pssm_file, 'rt') as ipf:
        for l in ipf:
            # skip empty lines or comment lines
            l_stripped = l.strip()
            if not l_stripped or l_stripped.startswith('#') or \
            l_stripped[0].isalpha():
                continue
            pssm_rows.append([float(x) for x in l_stripped.split()[2:-2]])
    return pssm_rows


def create_pssm_tensor(pssm, half_window_size=7):
    """
    

    Parameters
    ----------
    pssm : TYPE
        DESCRIPTION.
    half_window_size : TYPE, optional
        DESCRIPTION. The default is 7.

    Returns
    -------
    pssm_tensor : TYPE
        DESCRIPTION.

    """
    n_res = len(pssm)
    pssm_tensor = []
    for i in range(n_res):
        start_idx = i - half_window_size
        end_idx = i + half_window_size + 1
        if start_idx < 0:
            pssm_tensor.append(
                pssm[start_idx:] + pssm[:end_idx]
            )
        elif end_idx > n_res:
            pssm_tensor.append(
                pssm[start_idx:] + pssm[:end_idx-n_res]
            )
        else:
            pssm_tensor.append(
                pssm[start_idx:end_idx]
            )
    return pssm_tensor


def parse_cmd_args():
    """
    Parse command line arguments.

    Returns
    -------
    args : TYPE
        DESCRIPTION.

    """
    parser = ArgumentParser(description="")
    
    parser.add_argument(
        '--query', '-q', dest='query', type=str, required=True,
        help='''An amino acid sequence in FASTA format for which secondary
        structures and transmembrane topologies are to be predicted.'''
    )
    parser.add_argument(
        '--pssm', '-p', dest='pssm', type=str, required=False,
        help='''Position specific scoring matrix obtained from 
        running the BCL pssm application.'''
    )
    parser.add_argument(
        '--cnn-model', dest='cnn_model', type=str, required=True,
        help='''A multitasking deep CNN model taking PSSM as input 
        and trained to predict secondary structures and transmembrane
        topologies.'''
    )
    parser.add_argument(
        '--lstm-model', dest='lstm_model', type=str, required=True,
        help='''A LSTM model taking predictions from the CNN model as input 
        and trained to predict what protein class the sequence belongs to.'''
    )
    parser.add_argument(
        '--output-prefix', '-o', dest='output_prefix', type=str, required=False,
        default='out', help='''Prefix that will be prepended to the filenames
        of all output files.'''
    )
    
    args = parser.parse_args()
    return args


def main():
    # get command-line arguments
    cmd_args = parse_cmd_args()
    
    if cmd_args.pssm is None:
        # create a PSSM for the query sequence
        run_hhblits(cmd_args.query, cmd_args.prefix)
    
        # read the PSSM into a NumPy array
        if not os.path.exists(cmd_args.prefix + '.pssm'):
            print(
                'No position-specific-scoring matrix found! Please check the log file.'
            )
            sys.exit(1)
            pssm = read_pssm(cmd_args.prefix + '.pssm')
    else:
        pssm = read_pssm(cmd_args.pssm)
    
    # create tensors by taking windows from the PSSM array
    pssm_tensor = create_pssm_tensor(pssm, half_window_size=10)
    
    # load the cnn model
    cnn_model = models.load_model(cmd_args.cnn_model)
    
    # feature labels
    sse = ['H', 'E', 'C']
    loc = ['M', 'O']
    ori = ['L', 'P', 'O']
    top = ['>', '<', 'O']
    
    # make predictions
    sse_prob, loc_prob, ori_prob, top_prob = cnn_model.predict(pssm_tensor)
    
    pred_sses = ''.join([sse[i] for i in np.argmax(sse_prob, axis=1)])
    pred_locs = ''.join([loc[i] for i in np.argmax(loc_prob, axis=1)])
    pred_oris = ''.join([ori[i] for i in np.argmax(ori_prob, axis=1)])
    pred_tops = ''.join([top[i] for i in np.argmax(top_prob, axis=1)])
    
    combined_predictions = []
    for a, b, c, d in zip(pred_sses, pred_locs, pred_oris, pred_tops):
        combined_predictions.append(a + b + c + d)
    sentence = ' '.join(combined_predictions)

    # tokenize test texts
    tokenizer = Tokenizer(num_words=54)
    tokenizer.fit_on_texts(sentence)
    sequences = tokenizer.texts_to_sequences(sentence)
    X = pad_sequences(sequences, maxlen=1000)

    # load the LSTM model
    lstm_model = models.load_model(cmd_args.lstm_model)
    class_prob = lstm_model.predict(X)
    
    protein_classes = ['bitopic', 'tm-alpha', 'tm-beta', 'soluble']
    pred_classe = [protein_classes[i] for i in np.argmax(class_prob, axis=1)]
    
    # write predictions to disk files
    with open(cmd_args.prefix + '_predicted.sse', 'wt') as opf:
        opf.write(pred_sses)
        opf.write('\n')
    with open(cmd_args.prefix + '_predicted.loc', 'wt') as opf:
        opf.write(pred_locs)
        opf.write('\n')
    with open(cmd_args.prefix + '_predicted.ori', 'wt') as opf:
        opf.write(pred_oris)
        opf.write('\n')
    with open(cmd_args.prefix + '_predicted.top', 'wt') as opf:
        opf.write(pred_tops)
        opf.write('\n')
        
    # print a message
    print('The given sequence was predicted to be ' + pred_classe + '.')
    print('Predictions of individual features have been written to these files:')
    print(cmd_args.prefix + '_predicted.sse')
    print(cmd_args.prefix + '_predicted.loc')
    print(cmd_args.prefix + '_predicted.ori')
    print(cmd_args.prefix + '_predicted.top')
    print('Thanks for using MASSP!')


if __name__ == '__main__':
    main()
```
