## Supplementary material for "A multi-task deep-learning system for predicting membrane associations and secondary structures of proteins": Code and Raw Data: process_sse_prediction.py

```
import sys


ciphipsi_file = sys.argv[1]
top_file = sys.argv[2]
prefix = sys.argv[3]

# get the set of positions with missing residue
missing_res_set = set()
with open(ciphipsi_file, 'rt') as ipf:
    for l in ipf:
        if not l.startswith('#'):
            if 'X' in l or 'p' in l or 'a' in l:
                missing_res_set.add(int(l.split()[0]))
                
with open(top_file, 'rt') as ipf:
    header = next(ipf).strip()
    top_str = ''.join([l.strip() for l in ipf])

new_top_str = ''
for i, r in enumerate(top_str, start=1):
    if i in missing_res_set:
        continue
    else:
        new_top_str += r
        
with open(prefix + '_processed_sse.fasta', 'wt') as opf:
    opf.write(header)
    opf.write('\n')
    opf.write(new_top_str)
    opf.write('\n')
```
