## Supplementary material for "A multi-task deep-learning system for predicting membrane associations and secondary structures of proteins": Code and Raw Data: SOV_refine.pl

```
#!/usr/bin/env perl
#
# SOV_refine.pl
#########################################################################################
# The script is used to calculate SOV scores (SOV_refine and SOV'99)
#
# References: 
# (1) Tong Liu and Zheng Wang. SOV_refine: A further refined definition of 
# segment overlap score and its significance for protein structure similarity. 
# Source Code for Biology and Medicine, 2018, 13:1.
# (2) Zemla, A. et al. A modified definition of Sov, 
#            a segment-based measure for protein secondary
#            structure prediction assessment. (1999)
# 
# usage: perl SOV_refine.pl reference_sequence_file predicted_sequence_file lambda_value 
# Author: Tong Liu
#########################################################################################
use strict;
use warnings;
$| = 1;

if ($#ARGV < 1) {
    print  STDERR "Usage: $0 needs at least two parameters:
           (1) reference sequence file (fasta format);
           (2) predicted sequence file (fasta format);
           (3) lambda, (default 1).\n";
    exit 1;
}
my $file_reference = $ARGV[0];
my $file_predicted = $ARGV[1];
my $lambda = 1;
if ($ARGV[2]) {
    $lambda = $ARGV[2];
}
my ($seq_reference, $seq_predicted, $L1, $L2);
open IN, "$file_reference" or die "Error, cannot open the file $file_reference.";
while (my $line = <IN>) {
    chomp $line; 
    $line =~ s/^\s+//; 
    $line =~ s/\s+$//;
    if (substr($line, 0, 1) ne '>') {
        $seq_reference .= $line;
    }
}close IN;
open IN, "$file_predicted" or die "Error, cannot open the file $file_predicted.";
while (my $line = <IN>) {
    chomp $line; 
    $line =~ s/^\s+//; 
    $line =~ s/\s+$//;
    if (substr($line, 0, 1) ne '>') {
        $seq_predicted .= $line;
    }
}close IN;
$seq_reference = uc $seq_reference;
$seq_predicted = uc $seq_predicted;
$L1 = length($seq_reference);
$L2 = length($seq_predicted);
if ($L1 != $L2 || $L1 == 0 || $L2 == 0) {
    print STDERR "Error, the length of the two sequences isn't the same.\n";
    exit 1;
}
my @states;
for my $c (split //, $seq_reference) {
    if (! ($c ~~ @states)) {
        push(@states, $c);
    }
}
my $num_states = @states;
my $delta_all = delta_all_reference($lambda, $seq_reference, $num_states);

my (@SOVs_99, @SOVs_refine, @N);
for (my $i = 0; $i <= $#states; $i++) {
    my $state = $states[$i];
    my (%segments1, %segments2);
    parse_segments($seq_reference, \%segments1, $state);
    parse_segments($seq_predicted, \%segments2, $state);
    #######################################################################################
    # For each state, get the sets of S(i) and S'(i)
    # S(i):  the set of all the overlapped segment pairs (s1, s2)
    # S'(i): the set of all segment pairs (s1, s2) for which there is no overlapping segmen
    my (@SA, @SiA);
    for my $key1 (sort {$a<=>$b} keys %segments1) {
        my $i1 = $segments1{$key1}->[0];
        my $i2 = $segments1{$key1}->[1];
        my $tmp_num = keys %segments2;
        my $tmp_count = 0;
        for my $key2 (sort {$a<=>$b} keys %segments2) {  
            my $j1 = $segments2{$key2}->[0];
            my $j2 = $segments2{$key2}->[1];
            if ($j2 < $i1 || $i2 < $j1) { #the two segments no overlap
                $tmp_count++;
            } else { #$key1 belongs to SA
                my @tmp_SS; 
                push(@tmp_SS, $key1); 
                push(@tmp_SS, $key2);
                push(@SA, \@tmp_SS);
            }      
        }
        if ($tmp_count == $tmp_num) { #$key1 belongs to SiA
            push(@SiA, $key1);
        }
    }
    my ($len_SA, $len_SiA);
    $len_SA = $len_SiA = 0;
    my $n_sa = @SA;
    my $n_sia = @SiA;
    if (@SiA) {
        for (my $k = 0; $k < $n_sia; $k++) {
            $len_SiA += $segments1{$SiA[$k]}->[1] - $segments1{$SiA[$k]}->[0] + 1;
        }
    }
    if (@SA) {
        for (my $k = 0; $k < $n_sa; $k++) { 
            $len_SA += $segments1{$SA[$k][0]}->[1] - $segments1{$SA[$k][0]}->[0] + 1;
        }
    }
    $N[$i] = $len_SA + $len_SiA;
    my $tmp_sov_refine = 0;
		my $tmp_sov_99 = 0;
    if (@SA) {
        for (my $k = 0; $k < $n_sa; $k++) {
            my $tmp_len1 = $segments1{$SA[$k][0]}->[1] - $segments1{$SA[$k][0]}->[0] + 1;  
            my $tmp_len2 = $segments2{$SA[$k][1]}->[1] - $segments2{$SA[$k][1]}->[0] + 1; 
            my ($tmp_minov, $tmp_maxov);
            ($tmp_minov, $tmp_maxov) = get_minov_maxov($segments1{$SA[$k][0]}->[0], $segments1{$SA[$k][0]}->[1], $segments2{$SA[$k][1]}->[0], $segments2{$SA[$k][1]}->[1]);        
            my @tmp_mins;
            my $tmp_delta_refine = 0;
            my $tmp_delta_99 = 0;
            # bonus for SOV_99
            my ($t1, $t2, $t3, $t4);
            $t1 = $tmp_maxov - $tmp_minov;
            $t2 = $tmp_minov;
            $t3 = int $tmp_len1 / 2;         
            $t4 = int $tmp_len2 / 2;
            push(@tmp_mins, $t1);
            push(@tmp_mins, $t2);     
            push(@tmp_mins, $t3); 
            push(@tmp_mins, $t4);  
            $tmp_delta_99 = get_min(\@tmp_mins);
            # bonus for SOV_refine
            $tmp_delta_refine = $delta_all * ($tmp_len1 / $L1) * ($tmp_minov/$tmp_maxov);
            if ($tmp_delta_refine > ($tmp_maxov - $tmp_minov)) { 
                $tmp_delta_refine = $tmp_maxov - $tmp_minov; 
            }
            
            $tmp_sov_99 += ($tmp_minov + $tmp_delta_99) * $tmp_len1 / $tmp_maxov;
            $tmp_sov_refine += ($tmp_minov + $tmp_delta_refine) * $tmp_len1 / $tmp_maxov;
        }
        $SOVs_99[$i] = $tmp_sov_99 / $N[$i];
        $SOVs_refine[$i] = $tmp_sov_refine / $N[$i];
    } else {
        $SOVs_99[$i] = 0;
        $SOVs_refine[$i] = 0;
    }
}# End for each state

my ($N_all, $SOV_all_99, $SOV_all_refine);
$N_all = $SOV_all_99 = $SOV_all_refine = 0;
for (my $i = 0; $i < @states; $i++) {
    $N_all += $N[$i];
    $SOV_all_99 += ($SOVs_99[$i] * $N[$i]);
    $SOV_all_refine += ($SOVs_refine[$i] * $N[$i]);
} 
 
$SOV_all_99 /= $N_all;
$SOV_all_refine /= $N_all;

####################################
# output the Accuracy and SOV scores
my $accuracy = 0;
for (my $i = 0; $i < $L1; $i++) {
    if (substr($seq_reference, $i, 1) eq substr($seq_predicted, $i, 1)) {
        $accuracy++;
    }
} 
$accuracy /= $L1;
print "N_L\t$L1\n";
print "N_S\t$num_states\n";
print "Lambda\t$lambda\n";
for (my $i = 0; $i < @states; $i++) {
  my $tmp_sovi = sprintf "%0.3f", $SOVs_99[$i];
  print "SOV_99_i\t$states[$i]\t$tmp_sovi\n";
	$tmp_sovi = sprintf "%0.3f", $SOVs_refine[$i];
	print "SOV_refine_i\t$states[$i]\t$tmp_sovi\n";
}

$accuracy = sprintf "%0.3f", $accuracy;
$SOV_all_99 = sprintf "%0.3f", $SOV_all_99;
$SOV_all_refine = sprintf "%0.3f", $SOV_all_refine;

print "Accuracy\t$accuracy\n";
print "SOV_99\t$SOV_all_99\n";
print "SOV_refine\t$SOV_all_refine\n";

sub get_minov_maxov {
    my ($i1, $i2, $j1, $j2) = @_;
    $i1++; $i2++; $j1++; $j2++;
    my @tmp;
    my $minov = 0;
    my $maxov;
    push(@tmp, $i1);
    push(@tmp, $i2);
    push(@tmp, $j1);
    push(@tmp, $j2);
    my $min = get_min(\@tmp);
    my $max = get_max(\@tmp);
    $maxov = $max - $min + 1;
    for (my $i = $min; $i <= $max; $i++) {
        if ($i >= $i1 && $i <= $i2 && $i >= $j1 && $i <= $j2) {
            $minov++;
        }
    }
    return ($minov, $maxov);
}

sub get_min {
    my $arr = shift;
    my $min = $arr->[0];
    for (my $i = 1; $i < @{$arr}; $i++) {
        if ($arr->[$i] < $min) {
            $min = $arr->[$i];
        }
    }
    return $min;
}

sub get_max {
    my $arr = shift;
    my $max = $arr->[0];
    for (my $i = 1; $i < @{$arr}; $i++) {
        if ($arr->[$i] > $max) {
            $max = $arr->[$i];
        }
    }
    return $max;
}

sub parse_segments {
    my ($seq, $seg, $ss) = @_;
    my $tmp_len = length($seq);
    my $seg_num = 0;
    my ($start_index, $end_index);
    my $pre_ss = 0; #the previous state of the current position
    for (my $i = 0; $i < $tmp_len; $i++) {
        if (substr($seq, $i, 1) ne $ss) { #end a segment or no segment ahead
            if ($pre_ss) { #end a segment
                $end_index = ($i-1);
                $seg->{$seg_num}->[0] = $start_index;
                $seg->{$seg_num}->[1] = $end_index;
            }
            $pre_ss = 0;
        } else { #should belong to a segment or the first element initializes a segment
            if ($pre_ss == 0) { #initialize a segment
                $seg_num++;
                $start_index = $i;
            }
            if ($i == ($tmp_len-1)) {
                $end_index = $i;
                $seg->{$seg_num}->[0] = $start_index;
                $seg->{$seg_num}->[1] = $end_index;
            }
            $pre_ss = 1;       
        }
    }
}

sub delta_all_reference {
    my ($lambda, $seq, $num_states) = @_;
    my $L = length($seq);
    my $index = substr($seq, 0, 1);
    my $i = 1;
    my %hash;
    for my $s (split //, $seq) {
        if ($s eq $index) {
            $hash{$i}++;
        } else {
            $i++;
            $index = $s;
            $hash{$i}++;
        }
    }
    my $tmp_delta_all = 0;
    my $num_segs = keys %hash;
    for my $key (sort {$a<=>$b} keys %hash) {
        $tmp_delta_all += ($hash{$key} / $L) ** 2;
    }
    $tmp_delta_all = $lambda * $num_states / $tmp_delta_all;

    return $tmp_delta_all;
}

# End
```
