## Supplementary material for "A multi-task deep-learning system for predicting membrane associations and secondary structures of proteins": Code and Raw Data: train_cnn.py

```
#!/usr/bin/env python3

import os
import argparse
import random
import numpy as np
import seaborn as sns
import pandas as pd
import matplotlib.pyplot as plt
from tensorflow.keras import layers
from tensorflow.keras import Input
from tensorflow.keras import models
from tensorflow.keras import optimizers
from tensorflow.keras import callbacks


def one_hot(prot_sequence):
    """
    Encodes the given protein sequence according to one-hot encoding.

    Parameters
    ----------
    prot_sequence : str
        The amino acid sequence of a protein.

    Returns
    -------
    NumPy NDArray

    """
    aas = ['A', 'R', 'N', 'D', 'C', 'Q', 'E', 'G', 'H', 'I',
           'L', 'K', 'M', 'F', 'P', 'S', 'T', 'W', 'Y', 'V']
    aa_bin_vectors = []
    for aa in prot_sequence:
        aa_bin_vectors.append([
            1 if x == aa else 0 for x in aas
        ])
    return np.array(aa_bin_vectors, dtype=float)


def read_pssm(pssm_file):
    """
    Read position-specific-scoring matrix into a NumPy 2DArray.

    Parameters
    ----------
    pssm_file : str
        File containing the position-specific scoring matrix of a protein.

    Returns
    -------
    NumPy 2DArray

    """
    pssm_rows = []
    with open(pssm_file, 'rt') as ipf:
        for l in ipf:
            # skip empty lines or comment lines
            l_stripped = l.strip()
            if not l_stripped or l_stripped.startswith('#') or \
            l_stripped[0].isalpha():
                continue
            pssm_rows.append([float(x) for x in l_stripped.split()[2:-2]])
    return pssm_rows


def create_pssm_windows(pssm, half_window_size=7):
    """
    Format a window of the PSSM for training ConvNets.

    Parameters
    ----------
    pssm : NumPy 2DArray

    half_window_size : int
        Number of residues to include on each side of the central residue.

    Returns
    -------
    list
        A list of 2DArrays.

    """
    n_res = len(pssm)
    pssm_windows = []
    for i in range(n_res):
        start_idx = i - half_window_size
        end_idx = i + half_window_size + 1
        if start_idx < 0:
            pssm_windows.append(
                pssm[start_idx:] + pssm[:end_idx]
            )
        elif end_idx > n_res:
            pssm_windows.append(
                pssm[start_idx:] + pssm[:end_idx-n_res]
            )
        else:
            pssm_windows.append(
                pssm[start_idx:end_idx]
            )
    return pssm_windows


def read_ciphipsi(ciphipsi_file):
    """
    The .ciphipsi file for a given protein contains the target labels for all
    residues in the protein. This function parses the given .ciphipsi file into
    NumPy arrays for training the ConvNets.

    Parameters
    ----------
    ciphipsi_file : str
        A given .ciphipsi file.

    Returns
    -------
    NumPy Array.

    """
    # all considered target variables
    ss_types = ['H', 'E', 'C']
    localizations = ['M', 'S']
    orientations = ['L', 'P', '-']
    # '>' is up, '<' is down, '-' means soluble
    origins = ['>', '<', '-']

    # binary encoding vectors
    ss_bin_vectors = []
    loc_bin_vectors = []
    orient_bin_vectors = []
    orig_bin_vectors = []

    # iterate through all residues
    with open(ciphipsi_file, 'rt') as ipf:
        for l in ipf:
            l_stripped = l.strip()
            if l_stripped.startswith('#'):
                continue
            # get targets
            ss_type, localization, orientation, origin = l_stripped.split()[3:]
            # binary encoding
            ss_bin_vectors.append([
                1 if x == ss_type else 0 for x in ss_types
            ])
            loc_bin_vectors.append([
                1 if x == localization else 0 for x in localizations
            ])
            orient_bin_vectors.append([
                1 if x == orientation else 0 for x in orientations
            ])
            orig_bin_vectors.append([
                1 if x == origin else 0 for x in origins
            ])

    # return NumPy arrays
    return np.array(ss_bin_vectors, dtype=float), \
           np.array(loc_bin_vectors, dtype=float), \
           np.array(orient_bin_vectors, dtype=float), \
           np.array(orig_bin_vectors, dtype=float)


def create_dataset(data_path=None, data_file=None, half_window_size=10):
    """
    Convert raw training data into NumPy NDArrays so that they can be used as
    input to training Keras models.

    Parameters
    ----------
    data_path : str
        Path to the folder where training data is stored.

    data_file : str
        Path to the file containing the 5-letter IDs of proteins.

    half_window_size : int
        Number of residues to include at one side of the central residue.

    Returns
    -------
    NumPy NDArray

    """
    X = []
    y = []
    with open(data_file, 'rt') as ipf:
        pdb_chains = [l.strip() for l in ipf if not l.strip().startswith('#')]

    for chain in pdb_chains:
        pssm = read_pssm(
            pssm_file=os.path.join(data_path, 'pssm', chain + '.pssm')
        )
        pssm_windows = np.array(create_pssm_windows(pssm, half_window_size))
        ss, loc, orient, orig = read_ciphipsi(
            ciphipsi_file=os.path.join(data_path, 'ciphipsi', chain + '.ciphipsi')
        )
        # only keep residues for which all target attributes are labeled
        all_avail = np.all(
            np.vstack(
                (
                    np.any(ss, axis=1),
                    np.any(loc, axis=1),
                    np.any(orient, axis=1),
                    np.any(orig, axis=1)
                )
            ),
            axis=0
        )
        X.append(pssm_windows[all_avail])
        y.append(
            [ss[all_avail], loc[all_avail], orient[all_avail], orig[all_avail]]
        )
    return X, y


def plot_loss(history):
    """
    Function for plotting training and validation loss

    Parameters
    ----------
    history : History
        Keras History object containing the information about everything
        happened during training.

    Returns
    -------
    None

    """
    # set plotting parameters
    sns.set(font_scale=1.5)
    # get the History object
    history_dict = history.history
    # loss values
    train_loss = history_dict['loss']
    val_loss = history_dict['val_loss']
    # number of training iterations
    epochs = range(1, len(train_loss) + 1)
    # create a Pandas DataFrame object for plotting
    loss_df = pd.DataFrame(
        data={
            'epochs': list(epochs) + list(epochs),
            'loss': np.concatenate((train_loss, val_loss)),
            'group': ['Training loss'] * len(train_loss) + [
                'Validation loss'] * len(train_loss)
        }
    )
    fig, ax = plt.subplots()
    sns.scatterplot(x='epochs', y='loss', hue='group', data=loss_df)
    plt.xlabel('Epochs')
    plt.ylabel('Loss')
    plt.legend(loc='upper right')
    handles, labels = ax.get_legend_handles_labels()
    ax.legend(handles=handles[1:], labels=labels[1:])


def plot_acc(history):
    """
    Function for plotting training and validation accuracy

    Parameters
    ----------
    history : History
        A Keras History object containing everything happened during training.

    Returns
    -------
    None

    """
    # set plotting parameters
    sns.set(font_scale=1.5)
    # get the History object
    history_dict = history.history
    # accuracy values
    train_acc_arr = np.array([
        history_dict['sse_accuracy'],
        history_dict['loc_accuracy'],
        history_dict['orient_accuracy'],
        history_dict['top_accuracy']
    ])
    train_acc = np.mean(train_acc_arr, axis=0)
    val_acc_arr = np.array([
        history_dict['val_sse_accuracy'],
        history_dict['val_loc_accuracy'],
        history_dict['val_orient_accuracy'],
        history_dict['val_top_accuracy']
    ])
    val_acc = np.mean(val_acc_arr, axis=0)
    # number of training iterations
    epochs = range(1, len(train_acc) + 1)
    # create a Pandas DataFrame object for plotting
    acc_df = pd.DataFrame(
        data={
            'epochs': list(epochs) + list(epochs),
            'accuracy': np.concatenate((train_acc, val_acc)),
            'group': ['Training accuracy'] * len(train_acc) + [
                'Validation accuracy'] * len(val_acc)
        }
    )
    fig, ax = plt.subplots()
    sns.scatterplot(x='epochs', y='accuracy', hue='group', data=acc_df)
    plt.xlabel('Epochs')
    plt.ylabel('Accuracy')
    plt.legend(loc='upper left')
    handles, labels = ax.get_legend_handles_labels()
    ax.legend(handles=handles[1:], labels=labels[1:])


def train_underfit_model(data_path, train_ids, validation_ids):
    """
    Train a simple small model that underfits the data.

    Parameters
    ----------
    data_path : str
        Path to the folder where training data is located.
    train_ids : str
        Path to the file containing the 5-letter IDs of proteins in the
        training set.
    validation_ids : str
        Path to the file containing the 5-letter IDs of proteins in the
        validation set.

    Returns
    -------
    history : Keras History
        A History object, which has a member history, a dictionary containing
        data about everything that happened during training.

    """
    # create datasets for training models
    half_window_size = 3
    train_X, train_y = create_dataset(data_path, train_ids, half_window_size)
    validation_X, validation_y = create_dataset(
        data_path, validation_ids, half_window_size
    )

    # randomize the order of examples in training set and validation set
    n_proteins = len(train_X)
    shuffled_idx = list(range(n_proteins))
    random.shuffle(shuffled_idx)
    train_X_shuffled = [train_X[i] for i in shuffled_idx]
    train_y_sse = [train_y[i][0] for i in shuffled_idx]
    train_y_loc = [train_y[i][1] for i in shuffled_idx]
    train_y_orient = [train_y[i][2] for i in shuffled_idx]
    train_y_top = [train_y[i][3] for i in shuffled_idx]

    # convert to NumPy arrays
    train_X_shuffled = np.vstack(train_X_shuffled)[:, :, :, np.newaxis]
    train_y_sse = np.vstack(train_y_sse)
    train_y_loc = np.vstack(train_y_loc)
    train_y_orient = np.vstack(train_y_orient)
    train_y_top = np.vstack(train_y_top)

    # validation set
    n_vals = len(validation_X)
    val_X = np.vstack(validation_X)[:, :, :, np.newaxis]
    val_y_sse = np.vstack([validation_y[i][0] for i in range(n_vals)])
    val_y_loc = np.vstack([validation_y[i][1] for i in range(n_vals)])
    val_y_orient = np.vstack([validation_y[i][2] for i in range(n_vals)])
    val_y_top = np.vstack([validation_y[i][3] for i in range(n_vals)])

    val_y = {
        'sse': val_y_sse,
        'loc': val_y_loc,
        'orient': val_y_orient,
        'top': val_y_top
    }

    # instantiate a multi-output model
    input_shape = (half_window_size * 2 + 1, 20, 1)
    input_tensor = Input(shape=input_shape)
    x = layers.Conv2D(8, (3, 3), activation='relu')(input_tensor)
    x = layers.MaxPooling2D((2, 2))(x)
    x = layers.Flatten()(x)
    x = layers.Dense(8, activation='relu')(x)

    # heads
    sse_prediction = layers.Dense(3, activation='softmax', name='sse')(x)
    loc_prediction = layers.Dense(2, activation='softmax', name='loc')(x)
    orient_prediction = layers.Dense(3, activation='softmax', name='orient')(x)
    orig_prediction = layers.Dense(3, activation='softmax', name='top')(x)

    # multi-task model
    model = models.Model(
        input_tensor,
        [
            sse_prediction,
            loc_prediction,
            orient_prediction,
            orig_prediction
        ]
    )

    model.compile(
        optimizers=optimizers.Adam(lr=1e-4, decay=1e-4),
        loss=[
            'categorical_crossentropy',
            'categorical_crossentropy',
            'categorical_crossentropy',
            'categorical_crossentropy'
        ],
        metrics=[
            'accuracy',
            'accuracy',
            'accuracy',
            'accuracy'
        ]
    )

    model.summary()

    history = model.fit(
        train_X_shuffled,
        [
            train_y_sse,
            train_y_loc,
            train_y_orient,
            train_y_top
        ],
        epochs=20,
        batch_size=64,
        validation_data=(val_X, val_y)
    )
    return history


def train_overfit_model(data_path, train_ids, validation_ids):
    """
    Train a large model that is bound to overfit the data.

    Parameters
    ----------
    data_path : str
        Path to the folder where training data is located.
    train_ids : str
        Path to the file containing the 5-letter IDs of proteins in the
        training set.
    validation_ids : str
        Path to the file containing the 5-letter IDs of proteins in the
        validation set.

    Returns
    -------
    history : Keras History
        A History object, which has a member history, a dictionary containing
        data about everything that happened during training.

    """
    # increase window size to 15
    half_window_size = 15
    train_X, train_y = create_dataset(data_path, train_ids, half_window_size)
    validation_X, validation_y = create_dataset(
        data_path, validation_ids, half_window_size
    )

    # randomize the order of examples in training set and validation set
    import random
    n_proteins = len(train_X)
    shuffled_idx = list(range(n_proteins))
    random.shuffle(shuffled_idx)
    train_X_shuffled = [train_X[i] for i in shuffled_idx]
    train_y_sse = [train_y[i][0] for i in shuffled_idx]
    train_y_loc = [train_y[i][1] for i in shuffled_idx]
    train_y_orient = [train_y[i][2] for i in shuffled_idx]
    train_y_top = [train_y[i][3] for i in shuffled_idx]

    # convert to NumPy arrays
    train_X_shuffled = np.vstack(train_X_shuffled)[:, :, :, np.newaxis]
    train_y_sse = np.vstack(train_y_sse)
    train_y_loc = np.vstack(train_y_loc)
    train_y_orient = np.vstack(train_y_orient)
    train_y_top = np.vstack(train_y_top)

    # validation set
    n_vals = len(validation_X)
    val_X = np.vstack(validation_X)[:, :, :, np.newaxis]
    val_y_sse = np.vstack([validation_y[i][0] for i in range(n_vals)])
    val_y_loc = np.vstack([validation_y[i][1] for i in range(n_vals)])
    val_y_orient = np.vstack([validation_y[i][2] for i in range(n_vals)])
    val_y_top = np.vstack([validation_y[i][3] for i in range(n_vals)])

    val_y = {
        'sse': val_y_sse,
        'loc': val_y_loc,
        'orient': val_y_orient,
        'top': val_y_top
    }

    # instantiate a multi-output model
    input_shape = (31, 20, 1)
    input_tensor = Input(shape=input_shape)
    x = layers.Conv2D(16, (3, 3), activation='relu')(input_tensor)
    x = layers.Conv2D(32, (3, 3), activation='relu')(x)
    x = layers.Conv2D(64, (3, 3), activation='relu')(x)
    x = layers.MaxPooling2D((2, 2))(x)
    x = layers.Flatten()(x)
    x = layers.Dense(32, activation='relu')(x)
    x = layers.Dense(64, activation='relu')(x)

    # heads
    sse_prediction = layers.Dense(3, activation='softmax', name='sse')(x)
    loc_prediction = layers.Dense(2, activation='softmax', name='loc')(x)
    orient_prediction = layers.Dense(3, activation='softmax', name='orient')(x)
    orig_prediction = layers.Dense(3, activation='softmax', name='top')(x)

    # multi-task model
    model = models.Model(
        input_tensor,
        [
            sse_prediction,
            loc_prediction,
            orient_prediction,
            orig_prediction
        ]
    )

    model.compile(
        optimizers=optimizers.Adam(lr=1e-4, decay=1e-4),
        loss=[
            'categorical_crossentropy',
            'categorical_crossentropy',
            'categorical_crossentropy',
            'categorical_crossentropy'
        ],
        metrics=[
            'accuracy',
            'accuracy',
            'accuracy',
            'accuracy'
        ]
    )

    model.summary()

    history = model.fit(
        train_X_shuffled,
        [
            train_y_sse,
            train_y_loc,
            train_y_orient,
            train_y_top
        ],
        epochs=20,
        batch_size=64,
        validation_data=(val_X, val_y)
    )
    return history


def train_best_model(data_path, train_ids, validation_ids, model_path):
    """
    Train a model that has the best performance on the validation data.

    Parameters
    ----------
    data_path : str
        Path to the folder where training data is located.
    train_ids : str
        Path to the file containing the 5-letter IDs of proteins in the
        training set.
    validation_ids : str
        Path to the file containing the 5-letter IDs of proteins in the
        validation set.
    model_path : str
        Path to the folder in which the best model will be stored.

    Returns
    -------
    history : Keras History
        A History object, which has a member history, a dictionary containing
        data about everything that happened during training.

    """
    # set window size to 10
    half_window_size = 10
    train_X, train_y = create_dataset(data_path, train_ids, half_window_size)
    validation_X, validation_y = create_dataset(
        data_path, validation_ids, half_window_size
    )
    n_proteins = len(train_X)
    shuffled_idx = list(range(n_proteins))
    random.shuffle(shuffled_idx)
    train_X_shuffled = [train_X[i] for i in shuffled_idx]
    train_y_sse = [train_y[i][0] for i in shuffled_idx]
    train_y_loc = [train_y[i][1] for i in shuffled_idx]
    train_y_orient = [train_y[i][2] for i in shuffled_idx]
    train_y_orig = [train_y[i][3] for i in shuffled_idx]

    # convert to NumPy arrays
    train_X_shuffled = np.vstack(train_X_shuffled)[:, :, :, np.newaxis]
    train_y_sse = np.vstack(train_y_sse)
    train_y_loc = np.vstack(train_y_loc)
    train_y_orient = np.vstack(train_y_orient)
    train_y_orig = np.vstack(train_y_orig)

    #
    n_vals = len(validation_X)
    val_X = np.vstack(validation_X)[:, :, :, np.newaxis]
    val_y_sse = np.vstack([validation_y[i][0] for i in range(n_vals)])
    val_y_loc = np.vstack([validation_y[i][1] for i in range(n_vals)])
    val_y_orient = np.vstack([validation_y[i][2] for i in range(n_vals)])
    val_y_top = np.vstack([validation_y[i][3] for i in range(n_vals)])

    val_y = {
        'sse': val_y_sse,
        'loc': val_y_loc,
        'orient': val_y_orient,
        'top': val_y_top
    }

    # instantiate a multi-output model
    input_tensor = Input(shape=(21, 20, 1))
    x = layers.Conv2D(16, (3, 3), activation='relu')(input_tensor)
    x = layers.Conv2D(32, (3, 3), activation='relu')(x)
    x = layers.MaxPooling2D((2, 2))(x)
    x = layers.Flatten()(x)
    x = layers.Dropout(0.25)(x)
    x = layers.Dense(64, activation='relu')(x)

    # heads
    sse_prediction = layers.Dense(3, activation='softmax', name='sse')(x)
    loc_prediction = layers.Dense(2, activation='softmax', name='loc')(x)
    orient_prediction = layers.Dense(3, activation='softmax', name='orient')(x)
    orig_prediction = layers.Dense(3, activation='softmax', name='top')(x)

    # multi-task model
    model = models.Model(
        input_tensor,
        [
            sse_prediction,
            loc_prediction,
            orient_prediction,
            orig_prediction
        ]
    )

    callbacks_list = [
        callbacks.EarlyStopping(
            monitor='val_loss',
            patience=10
        ),
        callbacks.ModelCheckpoint(
            filepath=os.path.join(model_path, 'best_cnn_model.h5'),
            monitor='val_loss',
            save_best_only=True
        )
    ]

    model.compile(
        optimizer=optimizers.Adam(lr=1e-4, decay=1e-4),
        loss=[
            'categorical_crossentropy',
            'categorical_crossentropy',
            'categorical_crossentropy',
            'categorical_crossentropy'
        ],
        metrics=[
            'accuracy',
            'accuracy',
            'accuracy',
            'accuracy'
        ]
    )

    model.summary()

    # create combined set after tuning hyperparameters
    X = np.concatenate((train_X_shuffled, val_X))
    y_sse = np.concatenate((train_y_sse, val_y_sse))
    y_loc = np.concatenate((train_y_loc, val_y_loc))
    y_orient = np.concatenate((train_y_orient, val_y_orient))
    y_top = np.concatenate((train_y_orig, val_y_top))

    model.fit(
        X,
        [
            y_sse,
            y_loc,
            y_orient,
            y_top
        ],
        epochs=10,
        batch_size=64,
        alidation_data=(val_X, val_y),
        callbacks=callbacks_list
    )


def parse_cmd_arguments():
    """
    Parse command-line arguments.

    Returns
    -------
    ArgumentParser
        An ArgumentParser object containing the parsed command-line options.

    """
    parser = argparse.ArgumentParser(description='''
        
    ''')
    parser.add_argument(
        '-p', '--data-path', dest='data_path', type=str, required=True,
        help='Path to the directory where data files are located.'
    )
    parser.add_argument(
        '-t', '--training-ids', dest='training_ids', type=str, required=True,
        help='''A file containing the 5-letter PDB IDs of proteins in the 
        training set.'''
    )
    parser.add_argument(
        '-v', '--validation-ids', dest='validation_ids', type=str, required=True,
        help='''A file containing the 5-letter PDB IDs of proteins in the 
        test set.'''
    )
    return parser.parse_args()


def main():
    """

    Returns
    -------

    """
    # parse command-line arguments
    cmd_args = parse_cmd_arguments()

    # create datasets
    data_path = cmd_args.data_path
    train_id_file = cmd_args.training_ids
    validation_id_file = cmd_args.validation_ids
    test_id_file = cmd_args.test_ids

    train_X, train_y = create_dataset(data_path, train_id_file, 10)
    validation_X, validation_y = create_dataset(data_path, validation_id_file, 10)
    test_X, test_y = create_dataset(data_path, test_id_file, 10)

    # train an underfit model
    train_underfit_model(data_path, train_id_file, validation_id_file)

    # train an overfit model
    train_overfit_model(data_path, train_id_file, validation_id_file)

    # train best model
    train_best_model(data_path, train_id_file, validation_id_file)


if __name__ == '__main__':
    main()
```
