## Supplementary material for "A multi-task deep-learning system for predicting membrane associations and secondary structures of proteins": Code and Raw Data: train_lstm.py

```
#!/usr/bin/env python3

import os
import argparse
import numpy as np
from tensorflow.keras.preprocessing.sequence import pad_sequences
from tensorflow.keras.preprocessing.text import Tokenizer
from tensorflow.keras.layers import Dense, Embedding, LSTM
from tensorflow.keras.models import Sequential
from tensorflow.keras import optimizers, callbacks


def parse_cmd_arguments():
    """
    Parse command-line arguments.

    Returns
    -------
    ArgumentParser
        An ArgumentParser object containing the parsed command-line options.

    """
    parser = argparse.ArgumentParser(
        description='''This script can be run to train an LSTM model to 
        classify the protein type of the input amino acid sequence.'''
    )
    parser.add_argument(
        '--train-x', dest='train_x', type=str, required=True,
        help='Path of the file containing the training set raw data'
    )
    parser.add_argument(
        '--train-y', dest='train_y', type=str, required=True,
        help='Path of the file containing the training set labels.'
    )
    parser.add_argument(
        '--val-x', dest='val_x', type=str, required=True,
        help='Path of the file containing the validation set raw data.'
    )
    parser.add_argument(
        '--val-y', dest='val_y', type=str, required=True,
        help='Path of the file containing the validation set labels.'
    )
    parser.add_argument(
        '--model-path', dest='model_path', type=str, required=True,
        help='Path to the folder where the best models are stored.'
    )
    return parser.parse_args()


def main():
    """

    Returns
    -------

    """
    args = parse_cmd_arguments()

    protein_classes = ['bitopic', 'tm-alpha', 'tm-beta', 'soluble']

    with open(args.train_x, 'rt') as ipf:
        texts = [l.strip() for l in ipf]

    with open(args.train_y, 'rt') as ipf:
        labels = [l.strip() for l in ipf]

    labels = np.array([
        [1 if c == l else 0 for c in protein_classes]
        for l in labels
    ])

    with open(args.val_x, 'rt') as ipf:
        val_texts = [l.strip() for l in ipf]

    with open(args.val_y, 'rt') as ipf:
        val_labels = [l.strip() for l in ipf]

    val_labels = np.array([
        [1 if c == l else 0 for c in protein_classes]
        for l in val_labels
    ])

    max_features = 54
    maxlen = 1000
    batch_size = 4

    # tokenize training texts
    tokenizer = Tokenizer(num_words=max_features)
    tokenizer.fit_on_texts(texts)
    sequences = tokenizer.texts_to_sequences(texts)
    word_index = tokenizer.word_index
    print('Found %s unique tokens.' % len(word_index))
    data = pad_sequences(sequences, maxlen=maxlen)
    print('Shape of data tensor', data.shape)

    # tokenize validation texts
    val_tokenizer = Tokenizer(num_words=max_features)
    val_tokenizer.fit_on_texts(val_texts)
    val_sequences = val_tokenizer.texts_to_sequences(val_texts)
    x_val = pad_sequences(val_sequences, maxlen=maxlen)
    y_val = val_labels

    indices = np.arange(data.shape[0])
    np.random.shuffle(indices)
    x_train = data[indices]
    y_train = labels[indices]

    model = Sequential()
    model.add(Embedding(max_features, 32))
    model.add(LSTM(16))
    model.add(Dense(4, activation='softmax'))
    model.summary()

    callbacks_list = [
        callbacks.EarlyStopping(
            monitor='val_loss',
            patience=10
        ),
        callbacks.ModelCheckpoint(
            filepath=os.path.join(args.model_path, 'best_recurrent_model.h5'),
            monitor='val_loss',
            save_best_only=True
        )
    ]

    model.compile(
        optimizer=optimizers.Adam(lr=1e-4, decay=1e-4),
        loss='categorical_crossentropy', metrics=['acc'])
    model.fit(
        x_train,
        y_train, epochs=100,
        batch_size=batch_size,
        validation_data=(x_val, y_val),
        callbacks=callbacks_list
    )


if __name__ == '__main__':
    main()
```
